# Site-specific proximity labeling at single residue resolution for identification of protein partners *in vitro* and on cells

**DOI:** 10.1101/2023.07.27.550738

**Authors:** Thomas G. Bartholow, Paul Burroughs, Susanna K. Elledge, James R. Byrnes, Lisa L. Kirkemo, Virginia Garda, Kevin K. Leung, James A. Wells

**Affiliations:** Department of Pharmaceutical Chemistry, University of California San Francisco, San Francisco, California, 94158, USA; MIT, Cambridge, MA 02142; Interline Therapeutics, Brisbane, CA 94005; 3T Therapeutics, S. San Francisco, CA 94128; Department of Cellular & Molecular Pharmacology, University of California San Francisco, San Francisco, California, 94158, USA

## Abstract

The cell surface proteome, or surfaceome, is encoded by more than 4000 genes, but we are only beginning to understand the complexes they form. Rapid proximity labeling around specific membrane targets allows for capturing weak and transient interactions expected in the crowded and dynamic environment of the surfaceome. Recently, a high-resolution approach called μMap has been described (Geri, J. B., Oakley, J. V., Reyes-Robles, T., Wang, T., McCarver, S. J., White, C. H., Rodriguez-Rivera, F. P., Parker, D. L., Hett, E. C., Fadeyi, O. O., Oslund, R. C., and MacMillan, D. W. C. (2020) *Science 367*, 1091–1097) in which an iridium (Ir)-based photocatalyst is attached to a specific antibody to target labeling of neighbors utilizing light-activated generation of carbenes from diazirine compounds via Dexter Energy Transfer (DET). Here we studied and optimized the spatial resolution for the method using an oncoprotein complex between the antibody drug, trastuzumab (Traz), and its target HER2. A set of eight single site-specific Ir-catalytic centers were engineered into Traz to study intra- and inter-molecular labeling *in vitro* and on cells by mass spectrometry. From this structurally well-characterized complex we observed a maximum distance of ∼110 Å for labeling. Labeling occurred almost uniformly over the full range of amino acids, unlike the residue specific labeling of other techniques. To examine on cell labeling that is specific to HER2 as opposed to simply being on the membrane, we compared the labeling patterns for the eight Traz-catalyst species to random labeling of membrane proteins using a metabolically integrated fatty acid catalyst. Our results identified 20 high confidence HER2 neighbors, many novel, that were more than 6-fold enriched compared to the non-specific membrane tethered catalyst. These studies define distance labeling parameters from single-site catalysts placed directly on the membrane target of interest, and more accurately compare to non-specific labeling to identify membrane complexes with higher confidence.

## Introduction

The human cell surface proteome, or surfaceome, is encoded by one-fourth of the genome and represents the cellular interface for communication with other cells including signaling, cell nutrient transport, and cell-cell contact ^1–3^. The surfaceome is also the target of virtually all biologics and half of small molecule drug targets^4^. Thus, in recent years there has been a surge of mass spectrometry-based proteomics approaches to probe how the surfaceome remodels from health to disease, in terms of protein expression^2,5–7^, post-translational modifications (PTMs) such as glycosylation,^8–11^ and proteolysis^12^. However, these approaches do not interrogate the changes in the landscape and dynamics of protein complexes that initiate cell signaling.

In the past decade, the field of protein interactomics has made huge advances to identify stable protein complexes within the 3D environment of cells, principally with affinity tagged pull-down mass spectrometry^13–18^.The success of this approach requires high affinity partners for interactions to survive the pull-down work-up. However, affinity constants for reversable associations in the membrane are relaxed dramatically because membrane proteins in 2D have only three of six degrees of entropic freedom^19–21^; thus, transient complexes are unlikely to remain associated following simple affinity pull-downs and sample work-up.

To address this problem, the field of proximity labeling proteomics (PLP) has emerged as a way to take covalent snap-shots of proteins *in situ* by localized generation of reactive species containing a covalent biotin affinity handle for purification^22,23^. Peroxidase-based PLP, such as ascorbate peroxidase (APEX)^24^, have pioneered our ability to label the local proteomes of large structures like synapses^25^ and cilia^26^within minute time-scales. However, peroxidases generate hydroxy radicals are long-lived (t_1/2_ ∼0.1 msec) and estimated to diffuse large distances, up to 3000Å^27^. This labeling distance is more than ten-times the dimensions of typical binary protein complexes, and prone to capture bystander proteins in crowded surfaceome environments.

Chemical crosslinking is a powerful high-resolution alternative for structural characterization of strong complexes or dynamic protein states^28^, but necessitates close proximity of particular cross-linked residues (typically two lysines within about 15-30Å). Such cross-linking reactions are challenging to analyze from complex cellular samples, and so have been mostly limited to purified complexes^29^. Thus, there remains a gap for probing transient protein-protein interactions in complex cellular milieus, especially membrane proteomes.

In a significant step forward, the Oslund, Fadeyi, and MacMillan groups recently described a new higher resolution proximity labeling approach, called μMap^30^. μMap locally generates short half-life carbenes (t_1/2_ ∼ 2-4 nsec) from biotinylated diazirine compounds through illumination of a protein bound iridium catalyst and Dexter Energy Transfer (DET). Carbenes are known to insert into C-H bonds, among others, and are thus expected to indiscriminately label a broad spectrum of amino acid types in relatively short distances, relative to tyrosine radicals. μMap was shown capable of labeling cell surface complexes with much higher confidence than peroxidase-based catalysts due to the much shorter half-life of the carbene. The original μMap method placed 6-8 Ir-based catalysts by a random N-hydroxy succinamide (NHS) labeling of lysines onto a secondary antibody that binds a primary antibody directed to the membrane protein of interest. Using this secondary antibody approach, μMap was estimated to label sites within an average of ∼500-600Å from the secondary antibody DET catalyst ^30^.

We hypothesized that a site-specific Ir conjugation onto the primary antibody could give even higher resolution to the technique, and thus higher confidence to identify protein neighbors. Here, we tested this hypothesis using a Fab derived from trastuzumab (Traz) whose structure is known in complex with its drug target, HER2. Through our *in vitro* studies, we confirmed broad labeling amino acids at single amino acid resolution, and that labeling is rather uniform up to a maximum labeling distance of ∼120Å from the Ir-catalyst site. Also, to improve the confidence in partner-labeling on cells, we determined enriched partners relative to a non-specific membrane bound Ir-catalyst. We identified four times more high confidence partners than were found using the μMap method of NHS labeling onto a secondary antibody. More than a dozen new candidate interacting partners for HER2 were discovered with strong functional links to breast cancer. These studies serve to calibrate the distance dependence for carbene labeling at higher resolution and provide greater sensitivity and confidence for identifying membrane neighbors when referenced to a non-specific membrane bound catalyst on cells.

## Results

### Experimental Strategy

To systematically evaluate the performance of single-site photo-catalyst Fab probes, both *in vitro* and on cells, we chose the Traz-HER2 system. The Fab for Traz is structurally well-characterized in complex with HER2^31^, which allowed us to assess distances from Ir-photo-catalysts to specific labeled sites *in vitro*. We have previously shown the Fab for Traz is highly amenable to single site-specific labeling using engineered methionines and their specific reaction with oxaziridines for stable and selective bioconjugation^32^.

Importantly, Traz has minimal impact on the signaling of HER2^33^, and so Fab-catalysts should not significantly affect the signaling neighborhoods identified on cells.

We divided our studies into three parts (**Figure 1**). First, we interrogated intramolecular self-labeling *in vitro* by installing Ir-photo-catalysts at eight different sites in both the VH and CH-1 domains of heavy and light chain in the Fab. Here we sought to understand how the biotin-carbene labeling patterns varied within the Traz Fab, as well as the distance dependence from the attachment site of Ir photo-catalyst. Using high resolution mass spectrometry, we evaluated (i) the precise sites of intra-molecular carbene labeling within the Fab, (ii) the distribution of residue type labelled by quantitative mass spectrometry, and (iii) the distance dependence from catalyst site to labeled amino acid by inspection of the crystal structure (PDB: 1N8Z) (**Figure 1A**). Second, we determined intermolecular target labeling characteristics by analyzing labeling patterns from each of the Ir-photo-catalyst Fabs when bound to HER2 *in vitro*, and again determined the precise sites and distance dependence of the labeling for the complexes (**Figure 1B**). Finally, we determined the HER2 interactome using the eight different Fab-catalysts bound to HER2 on SKBR3 cells^34^. SKBR3 cells are a well-known breast cancer cell line that over expresses HER2 leading to oncogenic activation (**Figure 1C**). These on-cell analyses were benchmarked against a reference NHS labeled secondary antibody strategy, as typically done using μMap. Importantly, as a non-specific cellular control, we created a Ir-catalyst attached by click-chemistry to an azido-fatty acid (FA) that readily inserts into the cell membrane. This FA-catalyst control created an unbiased membrane proteomic background to which both primary and secondary antibody catalysts could be quantitatively compared for higher confidence interactome identification. We also show the FA-catalyst control can serve as a general approach for membrane PLP.

**Figure 1.**
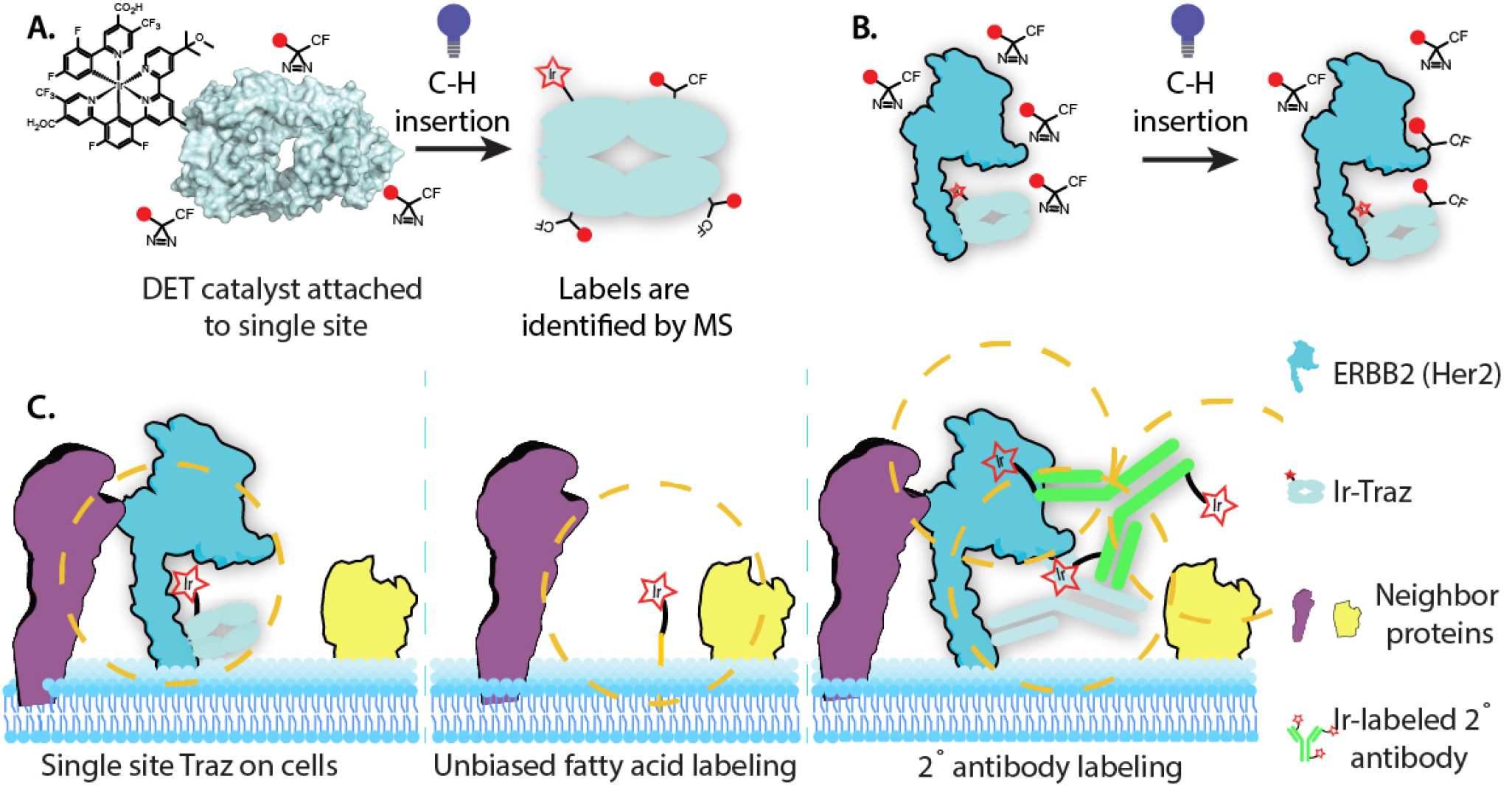
Experimental strategy to probe intramolecular, intermolecular, and on-cell biotin-carbene labeling by a Ir-based DET-catalysts that are site-specifically attached to the Traz Fab which binds HER2. **A**. Intramolecular biotin-carbene labeling for Ir-catalysts attached at single sites on the Traz-Fab. Labeling is catalyzed upon blue light illumination, which triggers localized biotin-carbene generation from the diazirine. Quantitative mass spectrometry was used to determine the precise sites labeled by biotin-carbenes, and the known structure (PDB:1N8Z) was used to assess distance dependence. **B**. Intermolecular labeling from the Fab-catalyst to HER2 in the complex was performed using same procedures as in **A**, to assess distance dependence from different Fab-catalyst probes. **C**. On-cell biotin-carbene labeling of cells expressing HER2 bound with specific Fab-catalysts (left) or secondary IgG catalyst (right) compared to the FA-catalyst non-specific control (control). HER2 is in blue and hypothetical partner in purple and by-stander in yellow.

### Analysis of intramolecular labeling of the Traz Fab from the Ir-photo catalyst positioned at eight sites

To create the Fab-catalysts, we expressed eight different single surface Met variants among 60 sites previously shown to be robustly expressed and quantitatively labeled with an oxaziridine for click chemistry^35^. The sites chosen were distributed throughout both light and heavy chains, in both the VH domain close to the CDRs as well as in the CH1 domain distal from the CDRs (**Supplemental Fig.1A)**. The eight Met Fab variants were expressed into the periplasm of *E. coli* and purified to >95%; each was labeled to >50% at the engineered single Met site using an oxaziridine-azide reagent via a sulfamide linkage; stoichiometry was confirmed by mass spectrometry (**Supplemental Fig. 2A**)^35^. The Traz Fab contains three buried Mets which have been shown not to react with the oxaziridine-azide reagent^35^. We next coupled the Ir-catalyst using copper-free click chemistry^36^; the final conjugated structure has a maximal extended linker length of ∼50Å (**Supplemental Fig 1B**).

We employed biotin-diazirine labeling using conditions similar to those for μMap^30^, where the Fab-catalyst concentration was set at 9uM, the biotin-diazirine concentration at 100uM (11-fold excess). Conjugates were illuminated with blue light for 5min at 25°C in phosphate buffered saline (PBS). The extent of labeling is controlled by the concentration of the biotin-diazirine, excess to protein, and time of illumination.

We then analyzed the sites of intramolecular biotin-carbene labeling in triplicate biological replicates each with two technical replicates for all eight Fab-catalysts at single amino acid resolution via mass spectrometry (**Figure 2**). Samples were digested with trypsin and peptides processed with a Preomics mass spectrometry prep kit^37^. Tryptic peptides were injected onto a Bruker TimsTOF pro and data processed using the PEAKS informatics program^38^. The location of the individual residues was readily identified by the presence of an additional monoisotopic mass of 616.25, corresponding to insertion of the biotin-carbene. Given the low sample complexity of tryptic peptides from these purified proteins, it was not necessary to enrich for the biotinylated peptides. Total coverage of peptides from the light chain was 98.6%, and for the heavy chain was 73.9%, when processed using a <1% FDR. While the coverage is very high, the tryptic peptides, S140-K153 and terminal S196-G239 of trastuzumab heavy chain were not detected.

**Figure 2.**
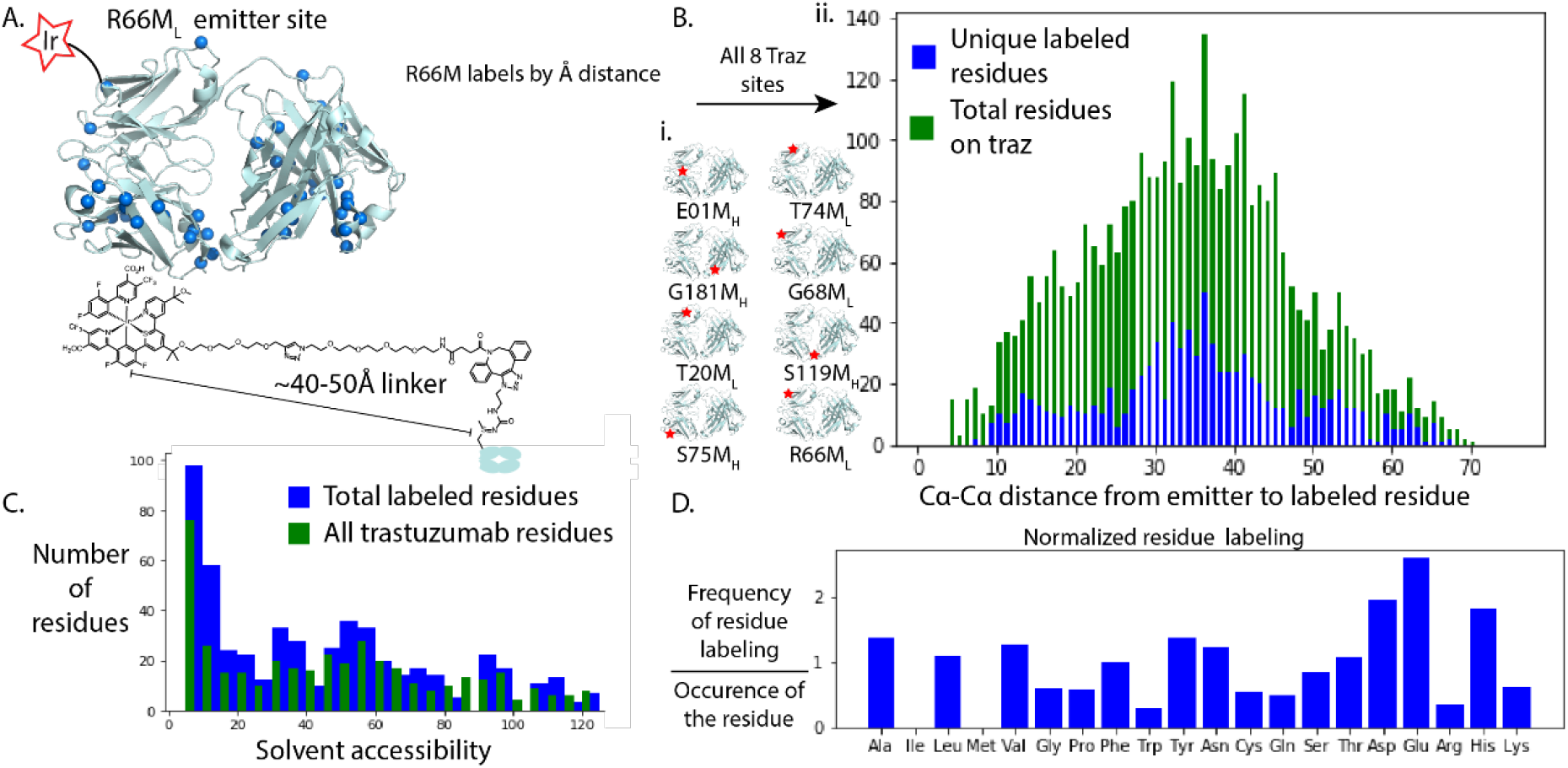
Dependence of distance, solvent accessibility, and residue type for intramolecular biotin-carbene labeling for eight Fab-catalyst probes. **A**. Ribbon diagram for a single Fab-catalyst (R66M_L_) where the Ir-catalyst was attached (red star). Locations of biotin-carbene labels are shown as blue dots. **B. i**. Ribbon diagrams for all eight Fab-catalysts with red stars showing the locations of the Ir-catalyst on the Fab, with specific mutation site indicated below. **ii**. Blue bars show a histogram for the Cα to Cα distance from the Ir-catalyst to sites of carbene labeling compiled for all eight Fab-catalysts. Note the gaussian shape centered at 30-40Å, and that labeling extends to the 65Å limit of the Fab’s longest dimension. Green bars show a histogram for Cα to Cα distance from the Ir-catalyst to all amino acids in the Fab **C**. The SASA value (calculated by Shrake-Rupley algorithm) for labeled residues (blue bars) and all residues (green bars). The ratio of green to blue is uniform, suggesting that labeling can occur if residue is partially or fully exposed. **D**. Normalized frequency of carbene labeling for the 20 amino acids, with a dotted line noting a 1:1 ratio. Although there is variation, most residues are uniformly labeled independent of amino acid type.

Each Fab-catalyst was covalently labeled with the biotin-containing probe at an average of 74 individual sites (+/-10), out of about 434 amino acid positions possible in the combined light and heavy chain. The precise sites of self-labeling were mapped onto the structure of the Fab-catalysts (**Figure 2A**; **Supplemental Figure 2C**). There was a large degree of overlap for the labeling patters for the eight Fab-catalysts although each was unique, as seen by the Venn diagrams in **Supplemental Figure 2B**.

From inspection of the crystal structure of Traz^31^, we determined the Cα to Cα distance from the methionine containing the Ir-catalyst to each biotin-carbene labeled amino acid for all from of the eight Fab-catalysts (blue bar graph shown in **Figure 2B)**. Summing the number of labels at each distance (based on a ∼500 total observations) produced a gaussian distribution centered around 30-40Å. We compared this to the a plot of the distances between any two residues (labeled or not) shown by the green bar graph overlayed in **Figure 2B**. This plot reflects the opportunity to label and shows a remarkably similar gaussian distribution centered around 30-40Å. If we then take the ratio of these two, for each distance position, this produced a rather flat plot that extends to the very edge of the Fab at 65Å. The decrease in labeling at the lower and higher angstrom distances matches the lower number of potential sites for labeling. We interpret this to mean that labeling potential is rather uniform to the limit of the Fab, ∼65Å.

We next analyzed the normalized frequency of labeling by amino acid type (**Figure 2D**). In aggregate, the labeling was broadly distributed across amino-acid types, and the labeling ratio varied only about +/-2-fold from the mean. There are ∼20 residues that are fully buried in the Fab, for which we did not see labeling, including all Ile and Met. We next determined how solvent accessible surface area (SASA) correlated with biotin-carbene labeling using standard Shrake-Rupley solvent accessibility calculations^39^ (**Figure 2C**). We found that carbene labeling could occur at all residues from partial to complete exposure (green bars), and this mirrored the average distribution of SASA for residues in the Fab (blue bars). Taken together, these data demonstrate that there is little bias for the residue type that is labeled, so long as they are partially solvent accessible, and that labeling is rather uniform to the edge of the Fab.

### Analysis of intermolecular labeling from Fab catalysts to HER2 in the complex

To evaluate the biotin-carbene labeling patterns for the Fab-catalysts in complex with the purified ecto-domain of HER2, we mixed them 1:1 (9uM each) and the conducted the biotin-carbene labeling. Given the high affinity of the Traz Fab for HER2 (K_D_ ∼ 2nM) ^40^ predicts that >95% of partners should be in 1:1 steady-state complex. Mass spectrometry workup and analysis was the same as for the Fab-catalyst self-labeling experiment above.

We evaluated how the self-labeling patterns for the Fab-catalysts changed when bound. The Fab-catalysts showed lower extents of self-labeling, with an average of 55 sites (+/-11) when bound to HER2 compared to an average of 74 sites (+/-10) when free. Some of this reduction may be because the HER2 complex is consuming available carbenes which is limiting. In addition, we see evidence that the CDRs can be partially protected from labeling when bound to HER2, as shown for the R66M_L_ Fab-catalyst in **Figure 3A**.

**Figure 3.**
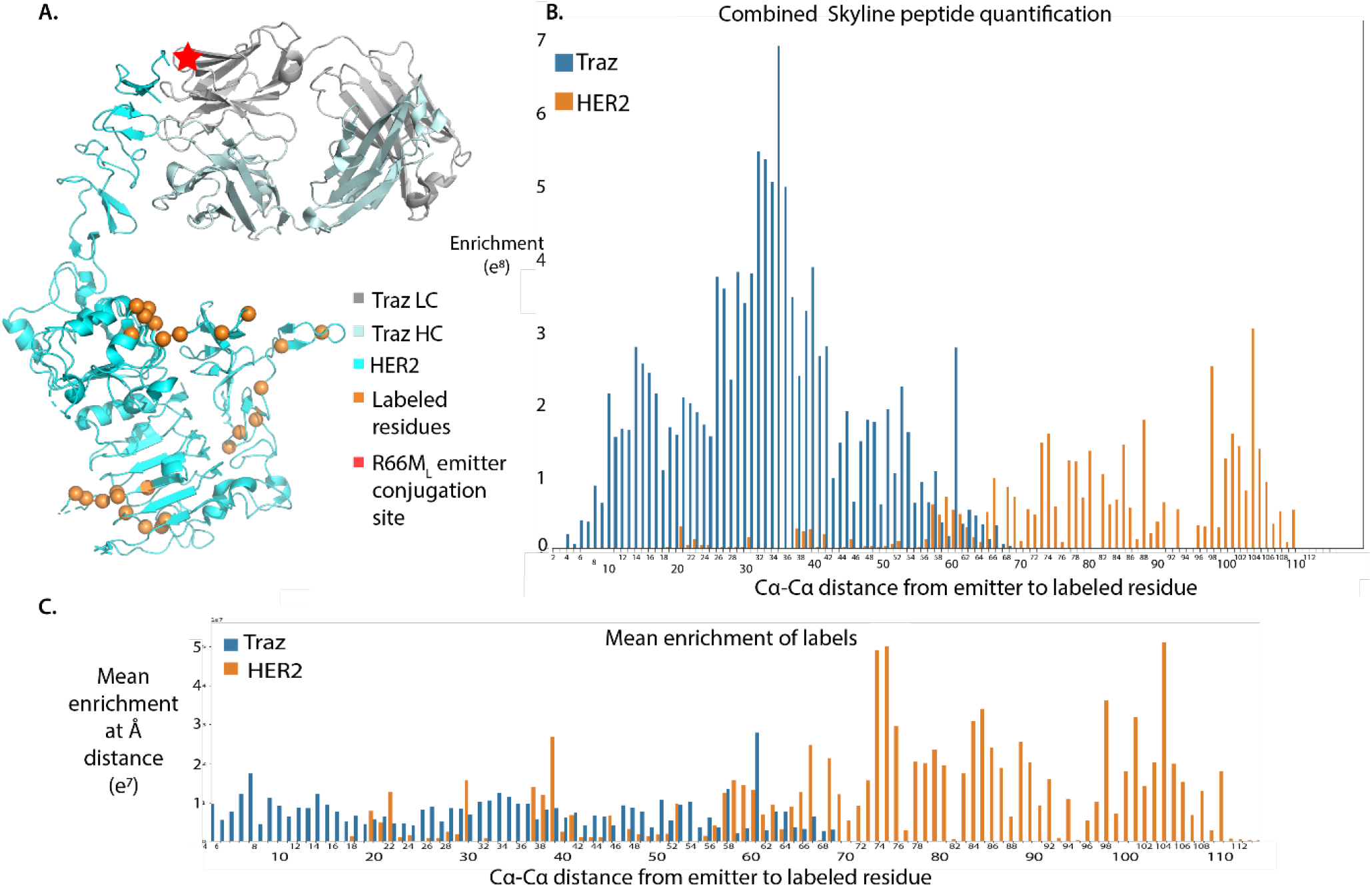
Intermolecular biotin-carbene labeling from Fab-catalyst in complex with HER2. **A**. Example of biotin-carbene labeling of HER2 using the R66M_L_ Fab catalyst. HER2 is rendered in cyan ribbons and Cα positions labeled are shown as orange spheres. The example R66M_L_ Fab-catalyst is shown in grey and teal ribbons, with the site of the Ir-catalyst shown as a red star. **B**. The total Cα to Cα distance distribution plot for intramolecular and intermolecular labeling, compiled for all eight Fab-HER2 complexes, showing the labeling of the Fab-catalysts (blue bars) and HER2 (orange bars). Labeling is graphed by enrichment calculated by Skyline. **C**. Enrichment of labeled residue at each distance from the Ir-catalyst taken as the average of enrichment at an angstrom distance.

We next analyzed the inter-molecular labeling patterns on HER2. The peptide coverage for HER2 averaged 52% with FDRs <1%; the “neck” region of HER2 is highly hydrophobic and poorly cleaved by Trypsin/LysC likely explaining why the peptides from this region were not resolved in the MS (red ribbons in **Figure 3B**). Each Fab-catalyst labeled between 11 to 28 sites on HER2, with an average of 16-sites (+/-12). The Fab-catalysts labelled many of the same sites on HER2 as shown by Venn diagrams in **Supplemental Fig 3A**,**B**, but each had a unique labeling pattern; this is not surprising given their different locations.

Next, we determined the distance dependence labeling for the complex, both intramolecular labeling of the Fab-catalyst alone, and intermolecular labeling of the Fab-catalyst complexed to HER2 (**Figure 3C**). For HER2, we see about ten times lower intensity of labeling than for the self-labeling by the Fab-catalyst. The labeling on HER2 extends from as close as ∼20Å from the closest Fab-catalyst to ∼110Å to the most distal. The furthest labeling distance possible on HER2 is 120Å.

We analyzed the extent of labeling versus distance from the Ir-catalyst for all eight complexes. For residues within ∼80Å of their most distant catalyst site, the extent labeling from different catalysts was very similar and generally flat (**Supplemental Figure 4**). However, for those sites with catalysts further than 100Å from the carbene labeled site, there was a sharp drop-off in extent of labeling for the more distal residues consistent with a maximum labeling near ∼110Å.

### Labeling of HER2 and its neighborhood on SKBR3 cells

We tested the ability of each of the eight Fab-catalysts to label HER2 and neighbors on a well-characterized breast cancer-derived cell line, SKBR3, where HER2 is over-expressed (1.6 million receptors per cell), leading to constitutive self-activation^34^. Each of the eight Fab-catalysts were added, at 5μg/mL (120nM), to 5 million SKBR3 cells for 30 min at 4°C. Excess Fab was quickly washed away followed by labeling with 100uM diazirine-biotin for 10 minutes of blue light exposure. We deliberately chose 4°C because it is below the lipid fluid phase transition so would effectively quench any further trafficking or diffusion. Cells were washed to remove labeling reagents, then lysed with RIPA. The biotinylated proteins were captured using neutravidin-agarose beads, digested on beads with trypsin, and eluted peptides analyzed by MS. Each Fab-catalyst sample was analyzed in biological triplicate and technical duplicate. Maxquant was used for processing the cell experiments, with Perseus used for further analysis. Details can be found in the Supplemental Methods. Each Fab-catalyst alone captured an average of 356 proteins (±21) **(Supplemental Table 3**). Together, the eight catalysts captured 742 proteins with >1 unique peptide per protein identified. About 90% of the captured proteins were annotated to be plasma membrane proteins based upon combination of SwissProt GOCC Plasma membrane and Uniprot databases (**Supplemental Fig. 5)**.

To rigorously compare proteins specifically labeled by the Fab-catalysts on the cell surface, we engineered a non-specific membrane catalyst through conjugation of the Ir-catalyst to a membrane anchored fatty acid. Five million SKBR3 cells containing the FA-catalyst were analyzed in biological triplicate with two technical replicates as before. The FA-catalyst identified a total of 787 proteins with high confidence (FDR < 1%) (**Supplemental Table 1**).

To further validate the FA-catalyst’s ability to act as an unbiased control to label the surfaceome, we compared its labeling pattern to the well-established method of cell surface capture (CSC). The CSC approach biotinylates the glycoprotein population of the cell surface, representing ∼85% of membrane proteins.^2^ We identified 543 proteins by CSC, with a standard deviation (SV) of 0.39 Log2x units between biologic replicates, close to the 0.32 SV value seen for the 787 proteins identified by the FA-catalyst control. There was substantial overlap with proteins identified by the FA-catalyst and CSC in both quantity and proteins identified (**Supplemental Fig 8 ; Supplemental Table 1**). Although there are some differences, it is likely these are due to the different cell surface labeling methods: the FA-catalyst labels anything within and expected 110Å and CSC is restricted to glycoproteins. Taken together, we believe this data validates the FA-catalyst for use for cell surfaceome labeling. Importantly, it allows us to compare methods (Fab vs. FA) that utilize the same Ir-based catalyst.

Next, we sought to compare our site-specific Fab-catalyst with the non-specific NHS labelling of the original uMap method ^30^. The uMap method used a mouse Fc secondary antibody labeled with roughly 6-8 Ir-catalysts. We produced this and tested labeling from solution as described in the μMap method. We identified only about 150 proteins, compared to 787 proteins for the FA-catalyst, with much less consistent LFQ values at a standard deviation of 0.67 (**Supplemental Table 1**). Virtually all of these were identified in the FA-catalyst control as shown by the VENN diagram (**Supplemental Fig. 5C**). These data support that the FA-catalyst more rigorously controls for non-specific membrane labeling of Fab-catalysts on the membrane than a solution borne antibody catalyst control.

We next analyzed labeling from the single-catalyst Fabs for the proteins that were enriched by proximity labeling. Remarkably, there was near complete overlap for the sum of all proteins detected at any level by each the eight Fab-catalysts and the FA-catalyst (**Figure 4Ai and 4Aii**). We quantitatively compared the protein-specific fold enrichment values for the each Fab-catalysts labeling versus the FA-catalyst control (**Figure 4B**). Proteins were filtered for those with at least 3 LFQ values in either one of the Fab-catalyst site mutants or in the FA-catalyst control. The result of this greater number of experiments was that there were more high confidence identifications, with 457 total proteins remaining in the dataset after the screening.

**Figure 4.**
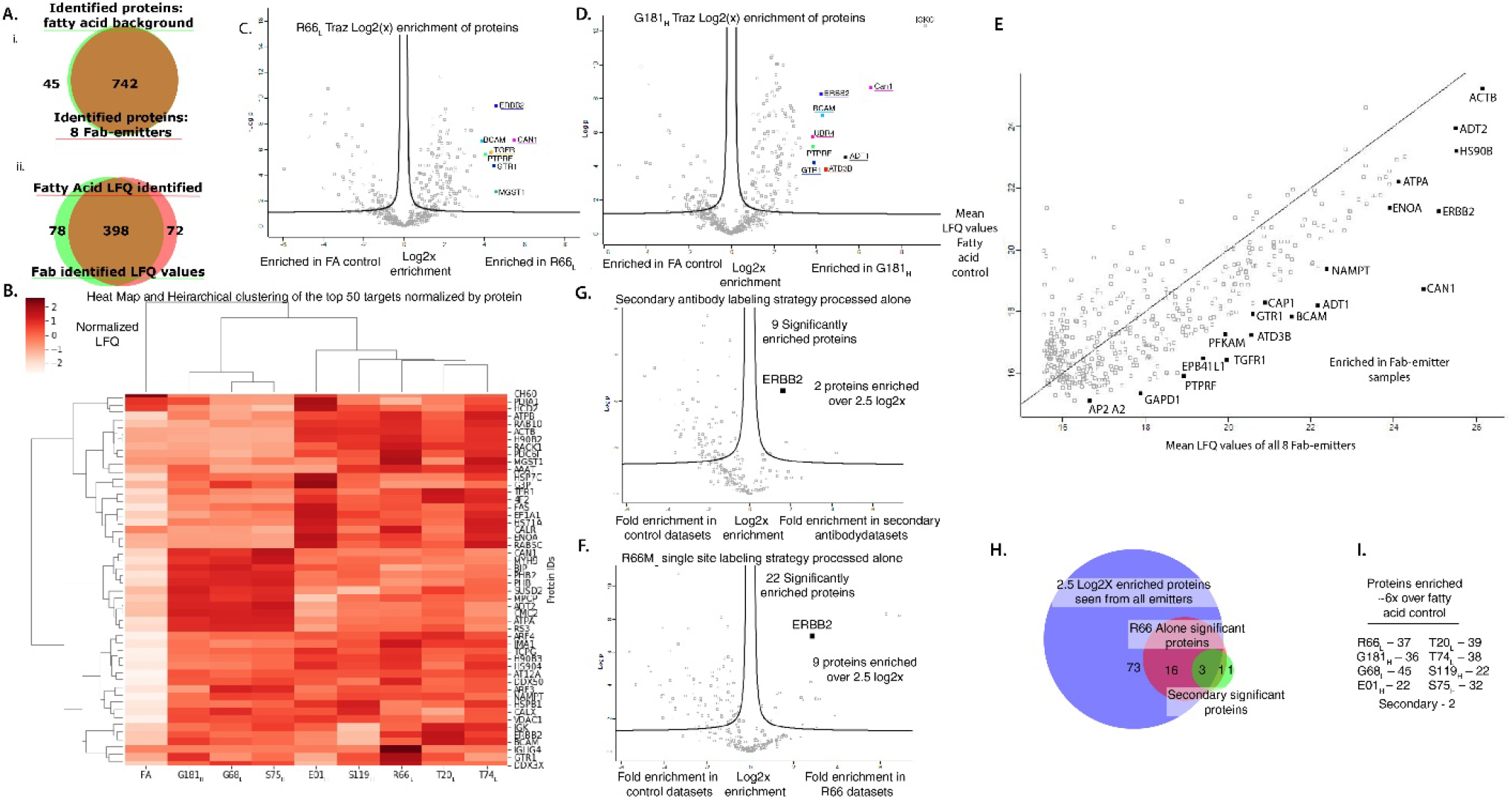
Proteins identified on SKBR3 cells for the eight Fab-catalysts and a secondary antibody-catalyst, compared to non-specific membrane labeling using the FA-catalyst. **A. i**. Venn diagram showing the overlap for all high confidence identified proteins identified in FA-catalyst and Fab-catalyst experiments. All high confidence proteins identified in the 8 Fab-catalysts were detected at some level in the FA-catalyst experiment. ii. Venn diagram showing the overlap of identified proteins with LFQ values. **B**. Non-hierarchical clustering and heat map of the top 50 most enriched proteins for the Fab-catalysts (lanes 2-9) relative to FA-catalyst (lane 1). LFQ values for each protein is normalized within the Fab samples. Non-normalized plots are available in **Supplemental Figure 8. C**. Volcano plot for R66M_L_ Fab-catalyst labeling processed together with all 8 Fab-catalysts. Proteins enriched greater than 2.5 log2 units or 6-fold over the FA-catalyst are labeled. **D**. G181M_H_ Fab-catalyst labeling processed identically to the R66M_L_, showing proteins enriched greater than 6-fold over the FA-catalyst. **E**. Mean enrichment of the eight Fab-catalysts plotted against the mean FA-catalyst enrichment levels. Proteins most enriched over FA-catalyst background are highlighted across a range of LFQ values. The overall range of LFQ values likely reflect differences in protein abundance. Proteins that fall furthest off the unity line are likely HER2 neighbors. Not surprisingly, the three of most highly enriched hits derive from HER2 and the Fab-catalyst light and heavy chains. **F**. Volcano plot showing fold enrichment for proteins labeled with the secondary antibody binding to Traz, relative to the FA-catalyst. Note there were only 9 high-confidence identifications, and only four that were enriched over 6-fold over the FA-catalyst control including HER2. **G**. A comparable volcano plot showing fold enrichment for proteins labeled with the R66M_L_ Fab-catalyst, compared against the FA-catalyst control; note there were 22 high-confidence identifications of which 9 proteins were enriched over 6-fold over the FA control. **H**. Venn diagram showing overlap of highly enriched proteins observed in all eight Fab-catalysts, compared to those in R66M_L_ only data set and the secondary antibody-catalyst dataset. **I**. List of eight Fab-catalysts and total number of proteins seen enriched for each by more than 6-fold relative to the FA-control.

Furthermore, a larger Maxquant run of 48 proteins identified more peptides for LFQ quantification, increasing the LFQ values with more total ID’s.

To compare the top 50 proteins with the highest enrichment values in the Fab-catalysts, we generated a heat map and nonhierarchical clustering (**Figure 4B**). Strikingly, the heat-map clearly distinguishes the group of eight Fab-catalysts from the FA-catalyst. In detail each Fab-catalyst is somewhat different, likely reflecting the different placement sites for the Ir-catalyst. These enrichment differences between Fab-catalyst and FA-catalysts can also be viewed by classic volcano plots (**Supplemental Figure 7**). To raise the statistical confidence of targets for each of the Fab-catalyst data sets, we combined them. This allowed less enriched hits to be retained relative to the FA-catalyst control. (**Figure 4C** and **D)** show example volcano plots for two of the Fab-catalysts, R66M_L_ and G181M_H_, which are furthest from each other on the Fab and show the greatest spread of differences.

Mean LFQ values from all Fab-catalysts were graphed against mean LFQ values for each FA-catalyst to determine enrichment (**Figure 4E**). Results demonstrate that across the entire range of LFQ values, specific proteins were selectively enriched by the Fab-catalysts against FA-control. The proteins labeled are those that are substantially and consistently enriched among the eight Fab-catalysts. As expected ERBB2 (HER2) and the primary antibody light and heavy chains from the Fab-catalysts are among the most abundant and enriched proteins.

We then sought to compare our site-specific labeled primary Fab-catalyst to the non-specific labeled secondary antibody approach described in uMap. We produced a comparable secondary antibody catalyst probe labeled with roughly 6-8 catalysts, which was then added to cells pre-bound with Traz. Samples were analyzed in biological triplicate and technical duplicate and processed as for the Fab-catalysts (**Supplemental Table 3**). In total, the secondary catalyst bound to Traz detected 334 proteins (<1% FDR), with 9 proteins significantly enriched over background and only five seen >6-fold enriched including the HER2 and the trastuzumab antibody (**Figure 4F; Supplemental Figure 6**). By comparison, we found 16 proteins enriched in the R66M_L_ Fab-catalyst by >6-fold, including virtually all those found in the targeted secondary antibody catalyst experiment (**Figure 4H**). These data indicate that site-specific catalyst attachment can provide higher resolution on cells than the multi-site NHS labeled secondary antibody catalyst.

Comparing volcano plots for all eight of the Fab-catalysts (**Supplemental Fig 7**), we see each Fab-catalyst has between 20 to 40 proteins enriched over 6-fold, relative to the FA-catalyst (**Figure 4H**). A Venn diagram of these highly enriched proteins shows each Fab-catalyst has a unique pattern (**Supplemental Figure 7)**, but we identify 16 proteins that are commonly labeled by four or more Fab-catalysts (**Supplemental Figure 7**).

Of the proteins consistently highly enriched, the majority are known to be secreted or membrane proteins, with 61 of 79 being assigned to the membrane in the GO database (**Supplemental Figure 7D)**. String analysis shows 71 of 88 proteins are known to have functional links to HER2 (**Supplemental Figure 6D**). We have further tabulated the common hits with the greatest amount of literature associated with HER2 biology or breast cancer (**Table I**).

**Table 1.** Selected proteins highlighted for cancer relevancy or HER2 signaling.

| MS ID | Protein name | Potentially relevance to Her2+ caner. |
| --- | --- | --- |
| CAN1 | Calpain-1 | A Protease linked to numerous disease states. Specifically tumor migration and progression<sup>50</sup>. |
| TGFR1 | TGF-beta receptor type-1 | A signaling protein linked to inhibition of early cancer progression, however in late cancer progression associated with increasing tumorigenicity and invasiveness<sup>51</sup>. |
| PTPRF | Receptor-type tyrosine-protein phosphatase F | A phosphatase which regulates signaling at cell-cell junctions. Overexpression has been implicated in decreased survivability in prostate cancer<sup>52</sup>. |
| BCAM | Basal cell adhesion molecule | An adhesion protein which through not fully established pathways has been linked to tumor progression, size, and negative survival rates<sup>53</sup>. |
| ENOA | Alpha-enolase | A glycolytic enzyme which is known to have a secondary role in cancers as a cell surface receptor increasing tumorigenicity<sup>54</sup>. |

## Discussion

### Single site in vitro experiments allow structure-based calibration of the carbene PLP labeling technology at single residue resolution

The recent development of μMap technology has opened exciting opportunities for higher resolution and higher confidence identification of protein partners on cells. In addition to better defining protein complexes *in situ*, we believe the technology has the theoretical potential to help provide structural constraints for protein complexes *in vitro*, much like spectroscopic methods like fluorescence energy transfer (FRET), or chemical cross-linking^28^. Thus, we focused on characterizing labeling distances and residues *in vitro* within a structurally well-defined complex at single residue resolution. This was accomplished by assessing sites of intramolecular carbene labeling for single Ir-catalysts placed at defined positions on the Traz Fabs, and sites of intermolecular labeling when bound to HER2. We quantified sites of carbene labeling at single amino-acid resolution from simple tryptic peptide mixtures without biotin enrichment.

From intra-molecular experiments on the site-specific Fab-catalysts, we found virtually all amino acids can be labelled, as anticipated from known chemistry of highly reactive carbenes^41^.

However, there were some systematic variations that were somewhat context dependent. We observed some biased labeling in favor of acidic residues, while noting a less than normalized labeling of basic residues. It is possible that this results from complementary and repulsive electrostatic interactions with the positively charged Ir-chelate. Such generally broad amino-acid coverage nicely allows labeling of many sites, thus increasing the sensitivity of detection. Other PLP methods that generate hydroxy radicals or reactive biotin-AMP react predominantly with tyrosine and lysines, respectively, are inherently less sensitive. About 20 highly buried residues were not labeled in the Traz-Fab, including Met and Ile. When correlated with SASA, we found that as long as an amino acid showed some exposure, it could be labeled.

With techniques to measure distances at the molecular scale, such as fluorescence energy transfer (FRET), nuclear overhauser effects (NOE) using NMR, and paramagnetic probes using EPR^42–44^, it is critical to empirically calibrate the distance dependence of effects on structurally defined sites with model systems. We applied the same reasoning to DET labeling here using intramolecular labeling of single-site defined Fabs and intermolecular labeling to its HER2 binding partner. Empirically, we see that labeling was rather uniform out to about 110Å. However, unlike FRET and NOE, we see considerable noise in the distance dependence of labeling. We hypothesize this has to do reaction efficiency, differences in how labeled peptides fly in the MS, and incomplete peptide coverage. Nonetheless, the broad and somewhat uniform amino acid coverage, the simplicity of MS analysis, and the availability of recombinant antibodies to many disease targets potentially can provide a useful orthogonal method to use for structural characterization even in complex samples.

The DET process involves an electron transfer from the singlet triplet excited state on the Ir-catalyst to the strained three-member ring of the diazirine within about 10 Å^45^. Once the carbene is triggered it can diffuse around 40 Å^46^ before it is quenched by water (given the half-life of the carbene in water of 7ns). In our system the Ir-catalyst is on a ∼50Å leash from the C? of the Met on the Fab-catalyst(estimated by Avogadro^47^). Summing the DET distance, diffusion, leash length we would theoretically expect a distance radius of labeling which is strongest at ∼70 Å and extreme distance of ∼110 Å possible which closely matches our measurements (**Figure 5**). Additional factors not captured here could include the dynamics of the leash and the protein, the potential binding of the Ir-chelate as it surveys the protein surface, and finally exclusion from penetrating the protein itself. All of these factors should affect sites that are labeled which is why it is critical to empirically determine distance relationships. Previous estimates using stimulated-emission depletion microscopy (STED)^27^ and three different constructs provided very useful distance ranges for biotin-carbene labeling for the μMap format^30^ of 570 +/-120 Å, 540 +/-120 Å, and 570+/-120 Å. If we overlay the estimates we have made upon the a primary and secondary antibody format which is detected by a streptavidin-fluorophore conjugate, a 60kDa tetramer of roughly 50Å diameter^30^ we can rationalize the ∼550Å range reported. Oakely et al^27^ also noted these effects would extend the range of the carbene labeling due to large size of these antibody-target-steptavidin complexes.

**Figure 5.**
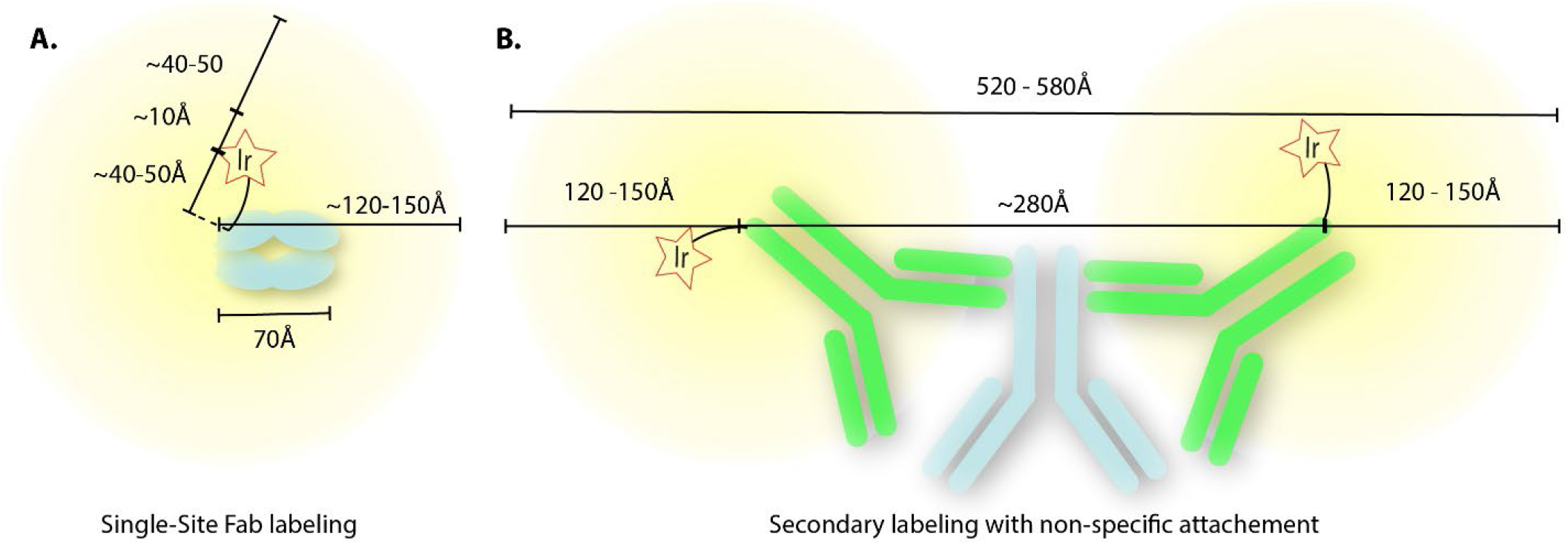
**A**. A cartoon schematic of the radial labeling distances we estimate compared to measurements. The theoretical distances are represented by the trio of determinants: a ∼10 Å distance for the DET process between the photocatalyst and diazirine, a ∼40-50 Å diffusion limit for carbenes in water, and a ∼40-50Å leash from the Ir-catalyst to the C? attachment site on the protein. These rougly match our maximum experimentally derived distances of ∼110-120 Å. **B** A schematic of this model applied to a secondary antibody system with approximate distances. This total distance agrees with the previously observed labeling distance of this complex of 540 ± 120 Å.

### On cell labeling with single-site Fab-catalysts revealed new members of signaling active HER2 neighborhoods with higher resolution and higher confidence

Trastuzumab is a small protein which does not perturb signaling. This allowed us to probe the active HER2 neighborhood in HER2 overexpressing cells, such as SKBR3 cells. We desired to establish if single site catalyst Fabs would perform better with their smaller radius of labeling compared to the μMap method, in terms of number and confidence of identifications and sensitivity for high and low abundance partners. To determine the specificity of labeling we hypothesized that a non-specific FA-tethered catalyst would more accurately replicate an unbiased sampling of the membrane protein background.

Membrane tethering of enzymes has shown to dramatically improve the depth of engaging membrane proteins by a sialydase^48^, a peroxidase^49^, or subtiligase^12^ by more than 20-fold. Thus, we created a general membrane anchored catalyst probe through attachment of the Ir-catalyst to a metabolically incorporated FA-catalyst. The FA-catalyst produced comparable numbers of identifications to traditional CSC methods for surfaceomics^8^ (between 600-1000 membrane proteins in a typical data set). By comparing the Fab-catalysts to the FA-catalyst, we identified about two dozen highly enriched partners (>6-fold) with high confidence for the Fab-catalyst probe (R66M_L_), while only four were labeled when the photo-catalyst was attached to using the secondary antibody μMap. We believe this reflects closer labeling from a single site primary Fab-catalysts that is positioned much closer to the HER2 target. The Fab-catalyst and FA-catalyst labeled largely the same ∼800 proteins on the cell but with differing LFQ values.

This is not surprising given the crowded nature and generally random distribution of proteins on the membrane. By bringing the photo-catalyst uniformly next to the target allowed better enrichment for those non-randomly associated proteins that interact with HER2.

All eight of the Fab-catalysts produced unique collections of two to three dozen high-confidence hits reflecting their different vantage points in labeling the neighborhoods. These are aggregate snap-shots - we cannot say that all these proteins form one large complex with HER2. Rather, we have identified proteins that were within ∼120Å for a longer period, specifically a set of common hits was found between all eight Fab-catalysts. Most of these have not been reported to bind with HER2 directly, but the majority have functional links to HER2 by STRING analysis. The identified proteins will require more detailed cellular experiments to validate functional biology with HER2. These could provide intriguing partner identifications for further study, and could provide useful for dual-targeting by bi-specific antibodies to give greater selectivity.

## Conclusions

The single-site Fab-catalyst probes extend the utility of high resolution carbene labeling. Here, we calibrated the method *in vitro* as a distance probe. Our studies demonstrate the broad breadth of residue-specific labeling and increased the resolution and sensitivity for site-specific attachment on the primary antibody. It is relatively simple to create single site catalysts to other binding proteins, which may be useful for assisting in characterizing structures and stoichiometries of protein complexes *in vitro* and in cells. Single-site catalyst probes allow one site to label surrounding proteins in a distance-restricted fashion. In addition, it is possible to increase and decrease the resolution by using catalysts with shorter or longer leashes or by switching out the reactive intermediate used. For example, recently it has been shown^27^ that photo-catalytic triggering of different phenyl azides to generate reactive nitrenes in uMap format increased labeling from 700Å to 1200Å, and with phenols up to ∼3000 Å. Given the abundance of recombinant proteins and antibodies, we believe single-site conjugated binding proteins, coupled with non-specific FA-catalyst controls, will increase the confidence and spatial resolution for transient protein interactomics.

## Acknowledgements

We are very grateful to Rob Oslund and Niyi Fadeyi for providing the Ir-photo catalyst and the biotin-diazrine substrate. We also thank them and colleagues Bernhard Geierstanger, Scott Lesley and Paul Marinec at Merck for very helpful discussions. We also thank Professor Andrej Sali and Dibyendu Mondal (UCSF) for useful structural considerations. The work was supported in part by a SEED grant from Merck. Additional funding for J.A.W. was from grants NIGMS R35GM122451, NCI R01CA248323, the CZ Biohub Investigator Program, the Harry and Dianna Hind Professorship.

## Notes

### Competing Interest Statement

The authors have declared no competing interest.

## References

(1) Carpenter, E. P., Beis, K., Cameron, A. D., and Iwata, S. (2008) Overcoming the challenges of membrane protein crystallography. Curr Opin Struct Biol 18, 581–586.

(2) Bausch-Fluck, D., Goldmann, U., Müller, S., van Oostrum, M., Müller, M., Schubert, O. T., and Wollscheid, B. (2018) The in silico human surfaceome. Proceedings of the National Academy of Sciences 115, E10988–E10997.

(3) Gonzalez, R., Jennings, L. L., Knuth, M., Orth, A. P., Klock, H. E., Ou, W., Feuerhelm, J., Hull, M. V., Koesema, E., Wang, Y., Zhang, J., Wu, C., Cho, C. Y., Su, A. I., Batalov, S., Chen, H., Johnson, K., Laffitte, B., Nguyen, D. G., Snyder, E. Y., Schultz, P. G., Harris, J. L., and Lesley, S. A. (2010) Screening the mammalian extracellular proteome for regulators of embryonic human stem cell pluripotency. Proceedings of the National Academy of Sciences 107, 3552–3557.

(4) Overington, J. P., Al-Lazikani, B., and Hopkins, A. L. (2006) How many drug targets are there? Nat Rev Drug Discov 5, 993–996.

(5) Bausch-Fluck, D., Hofmann, A., Bock, T., Frei, A. P., Cerciello, F., Jacobs, A., Moest, H., Omasits, U., Gundry, R. L., Yoon, C., Schiess, R., Schmidt, A., Mirkowska, P., Härtlová, A., Van Eyk, J. E., Bourquin, J.-P., Aebersold, R., Boheler, K. R., Zandstra, P., and Wollscheid, B. (2015) A mass spectrometric-derived cell surface protein atlas. PLoS One 10, e0121314.

(6) Wang, D., Eraslan, B., Wieland, T., Hallström, B., Hopf, T., Zolg, D. P., Zecha, J., Asplund Li, L., Meng, C., Frejno, M., Schmidt, T., Schnatbaum, K., Wilhelm, M., Ponten, F., Uhlen, M., Gagneur, J., Hahne, H., and Kuster, B. (2019) A deep proteome and transcriptome abundance atlas of 29 healthy human tissues. Molecular Systems Biology 15, e8503.

(7) Frei, A. P., Jeon, O.-Y., Kilcher, S., Moest, H., Henning, L. M., Jost, C., Plückthun, A., Mercer, J., Aebersold, R., Carreira, E. M., and Wollscheid, B. (2012) Direct identification of ligand-receptor interactions on living cells and tissues. Nat Biotechnol 30, 997–1001.

(8) Leung, K. K., Wilson, G. M., Kirkemo, L. L., Riley, N. M., Coon, J. J., and Wells, J. A. (2020) Broad and thematic remodeling of the surfaceome and glycoproteome on isogenic cells transformed with driving proliferative oncogenes. Proceedings of the National Academy of Sciences 117, 7764–7775.

(9) Fang, P., Ji, Y., Silbern, I., Doebele, C., Ninov, M., Lenz, C., Oellerich, T., Pan, K.-T., and Urlaub, H. (2020) A streamlined pipeline for multiplexed quantitative site-specific N-glycoproteomics. Nat Commun 11, 5268.

(10) Liu, M.-Q., Zeng, W.-F., Fang, P., Cao, W.-Q., Liu, C., Yan, G.-Q., Zhang, Y., Peng, C., Wu, J.-Q., Zhang, X.-J., Tu, H.-J., Chi, H., Sun, R.-X., Cao, Y., Dong, M.-Q., Jiang, B.-Y., Huang, J.-M., Shen, H.-L., Wong, C. C. L., He, S.-M., and Yang, P.-Y. (2017) pGlyco 2.0 enables precision N-glycoproteomics with comprehensive quality control and one-step mass spectrometry for intact glycopeptide identification. Nat Commun 8, 438.

(11) Riley, N. M., Hebert, A. S., Westphall, M. S., and Coon, J. J. (2019) Capturing site-specific heterogeneity with large-scale N-glycoproteome analysis. Nat Commun 10, 1311.

(12) Schaefer, K., Lui, I., Byrnes, J. R., Kang, E., Zhou, J., Weeks, A. M., and Wells, J. A. (2022) Direct Identification of Proteolytic Cleavages on Living Cells Using a Glycan-Tethered Peptide Ligase. ACS Cent. Sci. 8, 1447–1456.

(13) Babu, M., Krogan, N. J., Awrey, D. E., Emili, A., and Greenblatt, J. F. (2009) Systematic characterization of the protein interaction network and protein complexes in Saccharomyces cerevisiae using tandem affinity purification and mass spectrometry. Methods Mol Biol 548, 187–207.

(14) McGauran, G., Dorris, E., Borza, R., Morgan, N., Shields, D. C., Matallanas, D., Wilson, A. G., and O’Connell, D. J. (2020) Resolving the Interactome of the Human Macrophage aImmunometabolism Regulator (MACIR) with Enhanced Membrane Protein Preparation and Affinity Proteomics. PROTEOMICS 20, 2000062.

(15) Roux, K. J., Kim, D. I., Raida, M., and Burke, B. (2012) A promiscuous biotin ligase fusion protein identifies proximal and interacting proteins in mammalian cells. J Cell Biol 196, 801–810.

(16) Rhee, H.-W., Zou, P., Udeshi, N. D., Martell, J. D., Mootha, V. K., Carr, S. A., and Ting, A. Y. (2013) Proteomic mapping of mitochondria in living cells via spatially restricted enzymatic tagging. Science 339, 1328–1331.

(17) Richards, A. L., Eckhardt, M., and Krogan, N. J. (2021) Mass spectrometry-based protein– protein interaction networks for the study of human diseases. Molecular Systems Biology 17, e8792.

(18) Vermeulen, M., Hubner, N. C., and Mann, M. (2008) High confidence determination of specific protein–protein interactions using quantitative mass spectrometry. Current Opinion in Biotechnology 19, 331–337.

(19) Chataigner, L. M. P., Leloup, N., and Janssen, B. J. C. (2020) Structural Perspectives on Extracellular Recognition and Conformational Changes of Several Type-I Transmembrane Receptors. Frontiers in Molecular Biosciences 7.

(20) Sadeghi, M. (2022, January 25) Investigating the entropic nature of membrane-mediated interactions driving the aggregation of peripheral proteins. bioRxiv.

(21) Lindén, M., Sens, P., and Phillips, R. (2012) Entropic Tension in Crowded Membranes. PLoS Comput Biol 8, e1002431.

(22) Sears, R. M., May, D. G., and Roux, K. J. (2019) BioID as a Tool for Protein-Proximity Labeling in Living Cells. Methods Mol Biol 2012, 299–313.

(23) Deane, C. (2018) PUP up the volume. Nat Chem Biol 14, 903–903.

(24) Nguyen, T. M. T., Kim, J., Doan, T. T., Lee, M.-W., and Lee, M. (2020) APEX Proximity Labeling as a Versatile Tool for Biological Research. Biochemistry 59, 260–269.

(25) Loh, K. H., Stawski, P. S., Draycott, A. S., Udeshi, N. D., Lehrman, E. K., Wilton, D. K., Svinkina, T., Deerinck, T. J., Ellisman, M. H., Stevens, B., Carr, S. A., and Ting, A. Y. (2016) Proteomic analysis of unbounded cellular compartments: synaptic clefts. Cell 166, 1295.

(26) Mick, D. U., Rodrigues, R. B., Leib, R. D., Adams, C. M., Chien, A. S., Gygi, S. P., and Nachury, M. V. (2015) Proteomics of Primary Cilia by Proximity Labeling. Dev Cell 35, 497–512.

(27) Oakley, J. V., Buksh, B. F., Fernández, D. F., Oblinsky, D. G., Seath, C. P., Geri, J. B., Scholes, G. D., and MacMillan, D. W. C. (2022) Radius measurement via super-resolution microscopy enables the development of a variable radii proximity labeling platform. Proceedings of the National Academy of Sciences 119, e2203027119.

(28) Yu, C., and Huang, L. (2018) Cross-Linking Mass Spectrometry: An Emerging Technology for Interactomics and Structural Biology. Anal. Chem. 90, 144–165.

(29) Sinn, L. R., Giese, S. H., Stuiver, M., and Rappsilber, J. (2022) Leveraging Parameter Dependencies in High-Field Asymmetric Waveform Ion-Mobility Spectrometry and Size Exclusion Chromatography for Proteome-wide Cross-Linking Mass Spectrometry. Anal. Chem. 94, 4627–4634.

(30) Geri, J. B., Oakley, J. V., Reyes-Robles, T., Wang, T., McCarver, S. J., White, C. H., Rodriguez-Rivera, F. P., Parker, D. L., Hett, E. C., Fadeyi, O. O., Oslund, R. C., and MacMillan, D. W. C. (2020) Microenvironment mapping via Dexter energy transfer on immune cells. Science 367, 1091–1097.

(31) Cho, H.-S., Mason, K., Ramyar, K. X., Stanley, A. M., Gabelli, S. B., Denney, D. W., and Leahy, D. J. (2003) Structure of the extracellular region of HER2 alonez and in complex with the Herceptin Fab. Nature 421, 756–760.

(32) Elledge, S. K., Tran, H. L., Christian, A. H., Steri, V., Hann, B., Toste, F. D., Chang, C. J., and Wells, J. A. (2020) Systematic identification of engineered methionines and oxaziridines for efficient, stable, and site-specific antibody bioconjugation. Proceedings of the National Academy of Sciences 117, 5733–5740.

(33) Maadi, H., Nami, B., Tong, J., Li, G., and Wang, Z. (2018) The effects of trastuzumab on HER2-mediated cell signaling in CHO cells expressing human HER2. BMC Cancer 18, 238.

(34) Roberts, S. K., Hirsch, M., McStea, A., Zanetti-Domingues, L. C., Clarke, D. T., Claus, J., Parker, P. J., Wang, L., and Martin-Fernandez, A. M. L. (2018) Cluster Analysis of Endogenous HER2 and HER3 Receptors in SKBR3 Cells. Bio Protoc 8, e3096.

(35) Systematic identification of engineered methionines and oxaziridines for efficient, stable, and site-specific antibody bioconjugation - PubMed.

(36) Codelli, J. A., Baskin, J. M., Agard, N. J., and Bertozzi, C. R. (2008) Second-Generation Difluorinated Cyclooctynes for Copper-Free Click Chemistry. J. Am. Chem. Soc. 130, 11486–11493.

(37) Kulak, N. A., Pichler, G., Paron, I., Nagaraj, N., and Mann, M. (2014) Minimal, encapsulated proteomic-sample processing applied to copy-number estimation in eukaryotic cells. Nat Methods 11, 319–324.

(38) Tran, N. H., Qiao, R., Xin, L., Chen, X., Liu, C., Zhang, X., Shan, B., Ghodsi, A., and Li, M. (2019) Deep learning enables de novo peptide sequencing from data-independent-acquisition mass spectrometry. Nat Methods 16, 63–66.

(39) Shrake, A., and Rupley, J. A. (1973) Environment and exposure to solvent of protein atoms. Lysozyme and insulin. J Mol Biol 79, 351–371.

(40) Friedländer, E., Arndt-Jovin, D. J., Nagy, P., Jovin, T. M., Szöllősi, J., and Vereb, G. (2005) Signal transduction of erbB receptors in trastuzumab (Herceptin) sensitive and resistant cell lines: Local stimulation using magnetic microspheres as assessed by quantitative digital microscopy. Cytometry Part A 67A, 161–171.

(41) Doyle, M. P., Duffy, R., Ratnikov, M., and Zhou, L. (2010) Catalytic Carbene Insertion into C-H Bonds. Chem. Rev. 110, 704–724.

(42) Stryer, L., and Haugland, R. P. (1967) Energy transfer: a spectroscopic ruler. Proceedings of the National Academy of Sciences 58, 719–726.

(43) Kumar, A., Ernst, R. R., and Wüthrich, K. (1980) A two-dimensional nuclear Overhauser enhancement (2D NOE) experiment for the elucidation of complete proton-proton cross-relaxation networks in biological macromolecules. Biochem Biophys Res Commun 95, 1–6.

(44) Miao, Q., Nitsche, C., Orton, H., Overhand, M., Otting, G., and Ubbink, M. (2022) Paramagnetic Chemical Probes for Studying Biological Macromolecules. Chem. Rev. 122, 9571–9642.

(45) Arias-Rotondo, D. M., and McCusker, J. K. (2016) The photophysics of photoredox catalysis: a roadmap for catalyst design. Chem. Soc. Rev. 45, 5803–5820.

(46) Berg, H. C. (1993) Random Walks in Biology. Princeton University Press.

(47) Hanwell, M. D., Curtis, D. E., Lonie, D. C., Vandermeersch, T., Zurek, E., and Hutchison, G. R. (2012) Avogadro: an advanced semantic chemical editor, visualization, and analysis platform. Journal of Cheminformatics 4, 17.

(48) Gray, M. A., Stanczak, M. A., Mantuano, N. R., Xiao, H., Pijnenborg, J. F. A., Malaker, S., Miller, C. L., Weidenbacher, P. A., Tanzo, J. T., Ahn, G., Woods, E. C., Läubli, H., and Bertozzi, C. R. (2020) Targeted glycan degradation potentiates the anticancer immune response in vivo. Nat Chem Biol 16, 1376–1384.

(49) Kirkemo, L. L., Elledge, S. K., Yang, J., Byrnes, J. R., Glasgow, J. E., Blelloch, R., and Wells, J. A. (2022) Cell-surface tethered promiscuous biotinylators enable comparative smallscale surface proteomic analysis of human extracellular vesicles and cells. Elife 11, e73982.

(50) Storr, S. J., Carragher, N. O., Frame, M. C., Parr, T., and Martin, S. G. (2011) The calpain system and cancer. Nat Rev Cancer 11, 364–374.

(51) Syed, V. (2016) TGF-β Signaling in Cancer. J Cell Biochem 117, 1279–1287.

(52) Du, Y., and Grandis, J. R. (2015) Receptor-type protein tyrosine phosphatases in cancer. Chin J Cancer 34, 61–69.

(53) Chang, H.-Y., Chang, H.-M., Wu, T.-J., Chaing, C.-Y., Tzai, T.-S., Cheng, H.-L., Raghavaraju, G., Chow, N.-H., and Liu, H.-S. (2017) The role of Lutheran/basal cell adhesion molecule in human bladder carcinogenesis. Journal of Biomedical Science 24, 61.

(54) Almaguel, F. A., Sanchez, T. W., Ortiz-Hernandez, G. L., and Casiano, C. A. (2020) Alpha-Enolase: Emerging Tumor-Associated Antigen, Cancer Biomarker, and Oncotherapeutic Target. Frontiers in Genetics 11.

