## Supplemental figures and supplemental tables 1&2 for "Site-specific proximity labeling at single residue resolution for identification of protein partners *in vitro* and on cells"

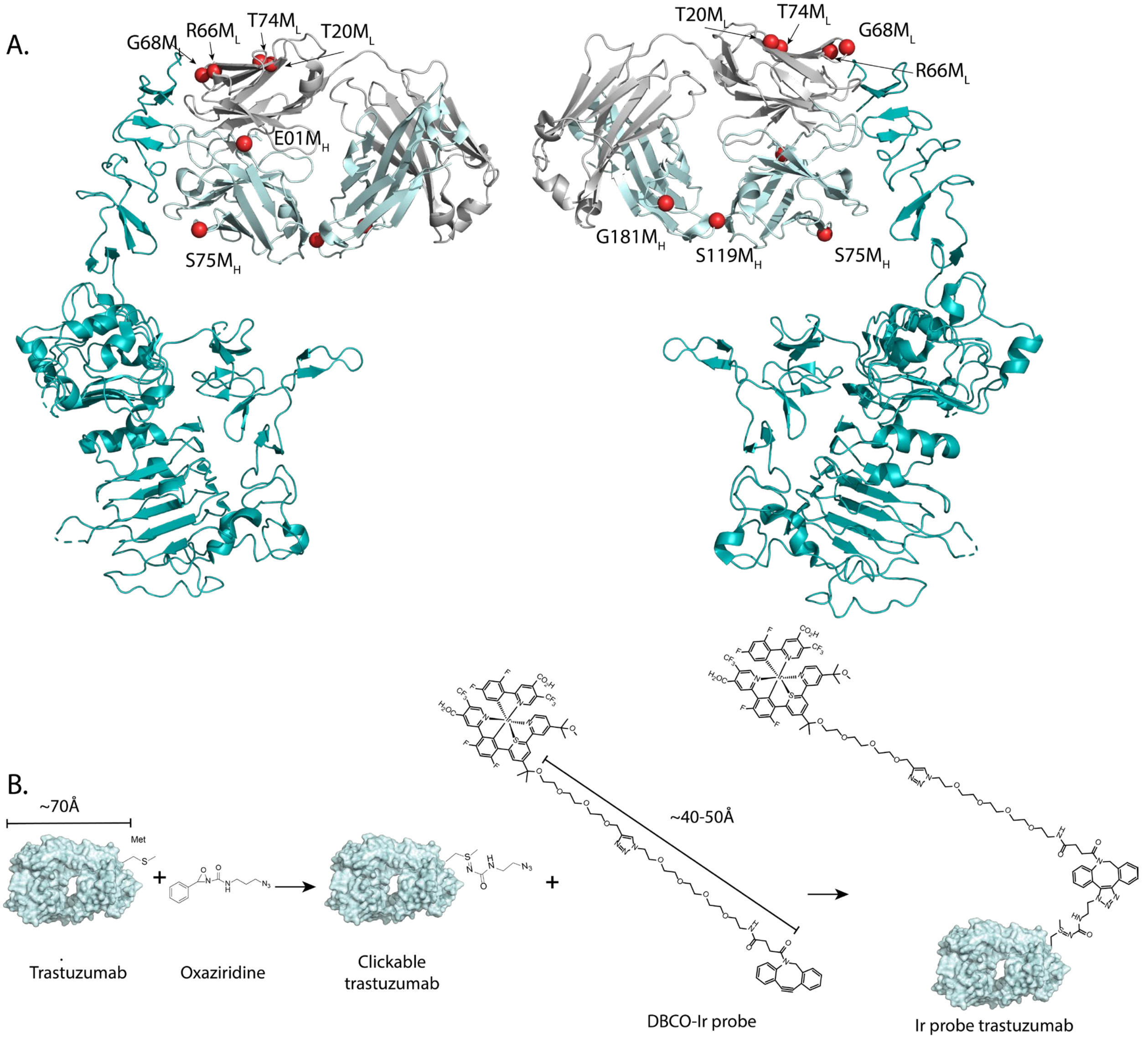


**SI Figure 1.** Methionine sites and conjugation chemistry used for step-wise addition of Ir-catalyst to the Fab of Traz. **A.** Ribbon diagram showing front and back view of the complex between the extracellular domain of HER2 bound with Traz. The eight sites across light and heavy chains in Traz that were mutated to methionine in order to perform site-specific oxaziridine conjugation are shown in red dots distributed. The HER2 structure is shown in Teal with the Traz light chain in gray and heavy chain in light cyan **B.** The two reaction steps involved to create the DET-probe conjugated to Traz are shown (i) oxaziridine conjugation chemistry onto the Met to attach the azide and (ii) attachment of the Ir catalyst using DBCO click chemistry. The protein is minimized in scale in order to highlight the structure of the probe.

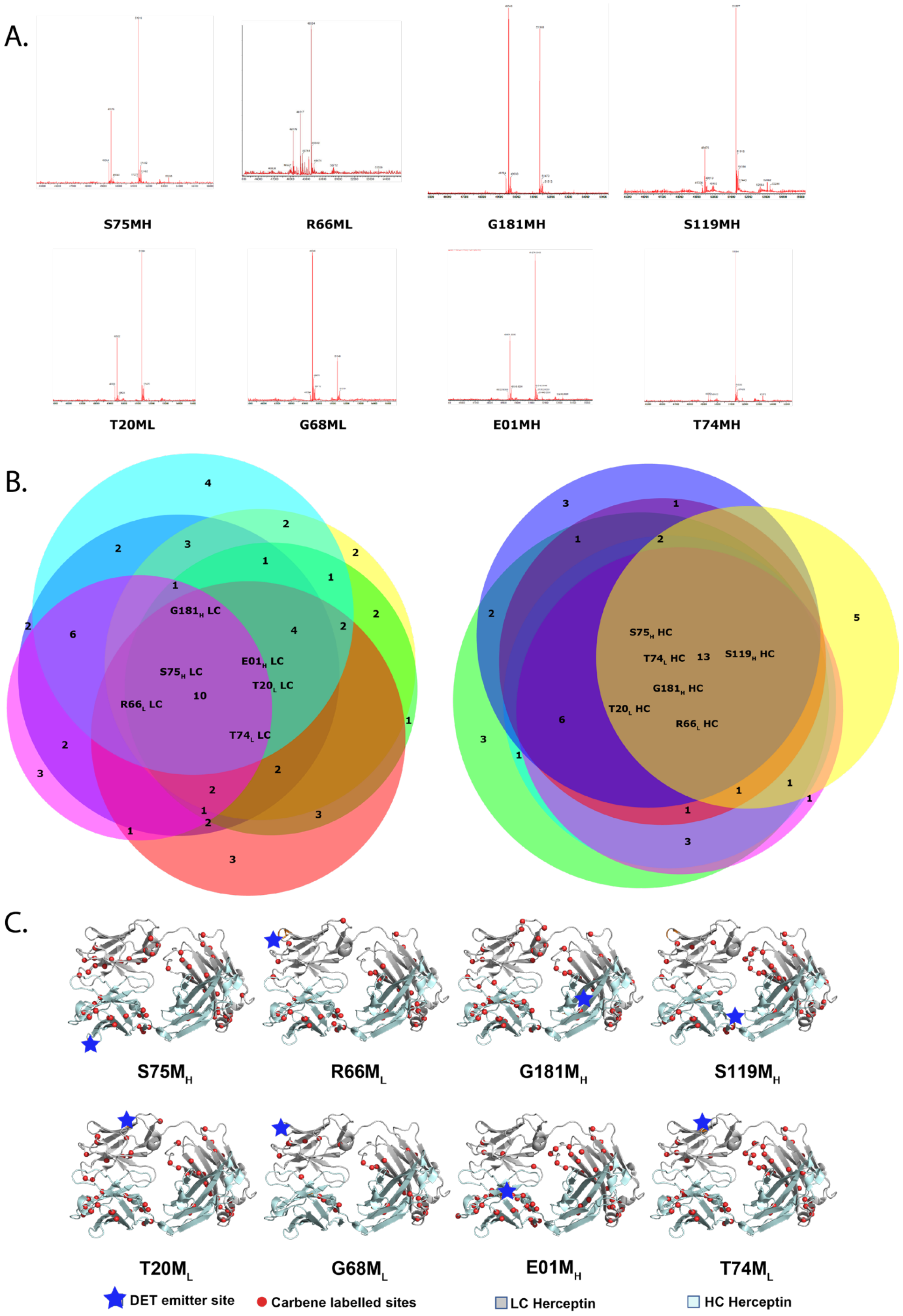


**SI Figure 2**. Intramolecular labeling of Traz by the Ir-catalyst from eight different Met sites. **A.** MS traces of the Ir-catalyst conjugated Fabs. 100% labeling was not achieved in all cases but was not found to be essential to experiments.**B.** Venn diagram of the labeling of for six of the eight catalyst sites on the Traz light and heavy chain (Light cyan and grey ribbons, respectively). Full reports of labeled sites are available in the supplemental tables. Most sites overlap but each catalyst has a unique detailed labeling pattern. **C.** Sites of carbene insertion (red dots) from each of the eight catalyst sites (shown by star).

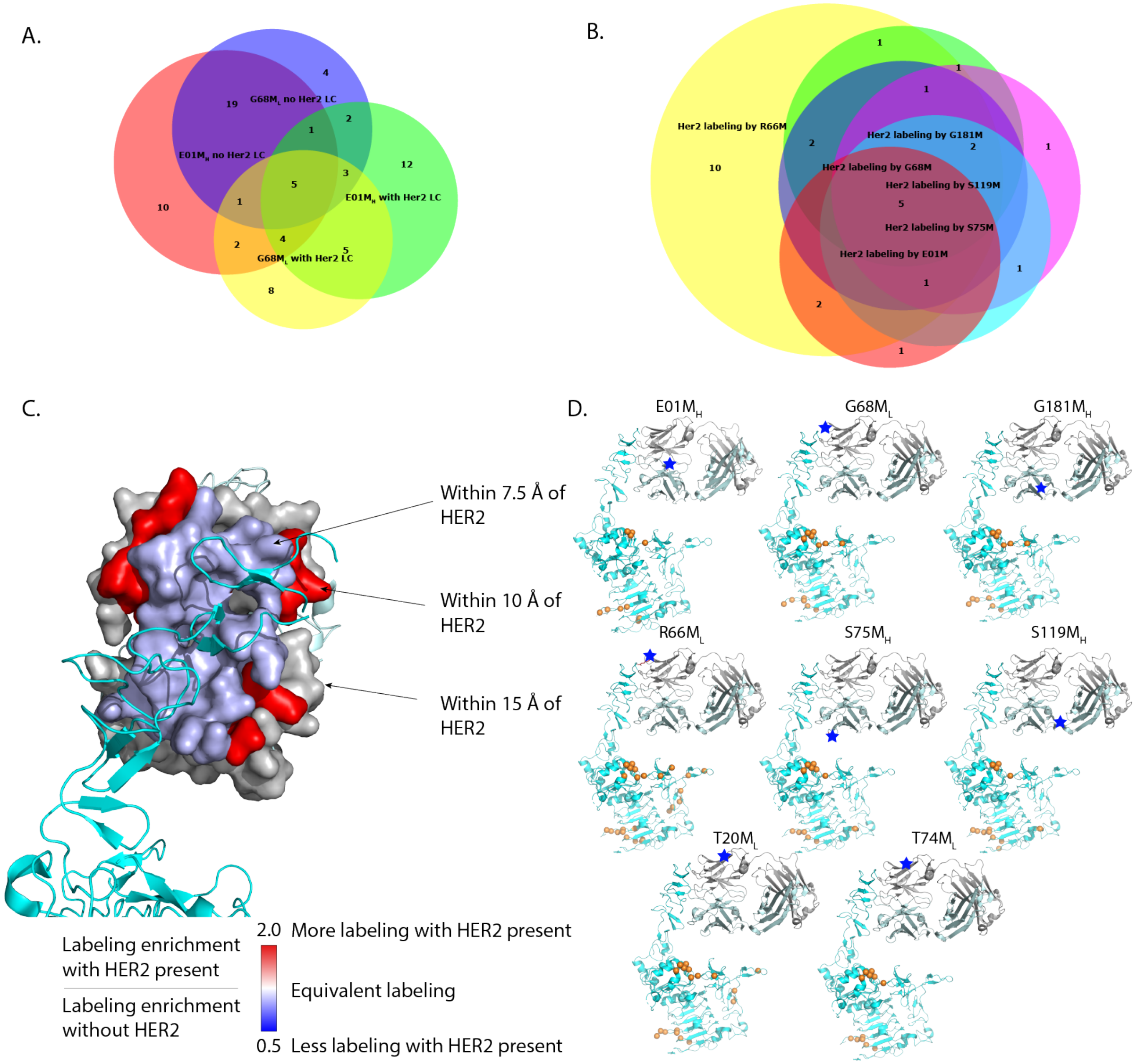


**SI Figure 3.** Carbene labeling of the HER2-Traz complex from each of the Traz Ir-catalyst conjugates **A.** Venn diagram of labeling comparing two emitter sites with and without HER2. Demonstrating that addition of the partner appears to have effects on where labels are seen. **B.** A Venn diagram of labeling by 6 of the 8 Fab-catalysts. Many of the carbene labeling sites are in common on HER2 although the closest one, R66M$_L$, systematically had more than the others. **C.** Assessment of the protection by HER2 of the emitter-Fab self-labeling. The contact surface of the emitter-Fab was rendered in space filling view and colored based on the ratio of labeling in the presence and absence of HER2 (teal ribbon). Ratios are generated by combining all the labeling seen for residues at the described distance from HER2. **D.** The sites of carbene labels (organ spheres on alpha carbons) seen from each of the eight catalysts. The HER2 structure is shown in Teal with the Traz light chain in gray and heavy chain in light cyan. Emitter sites are highlighted as blue stars.

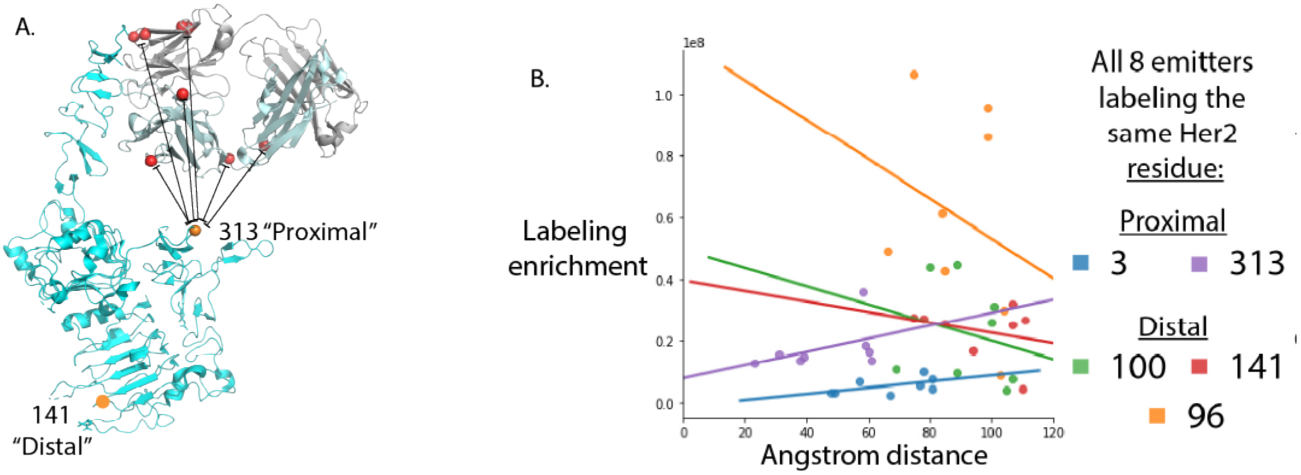


**SI Figure 4.** Distance dependence for the labeling of the same residue on HER2 from each of the eight catalyst sites on Traz. **A.** Structure of Traz (light cyan and grey ribbons) bound to HER2 (teal ribbons). Traz catalyst sites are indicated in red dots and example common proximal or distal residues on HER2 labeled by the carbene indicated in orange dots. **B.** Distance dependence for carbene labeling at five common residues on HER2 from the eight different catalysts on Traz. Note that proximal residues show relatively flat even profiles across distance while those distal show a loss at greater distance dependence and both terminate near 110Å.

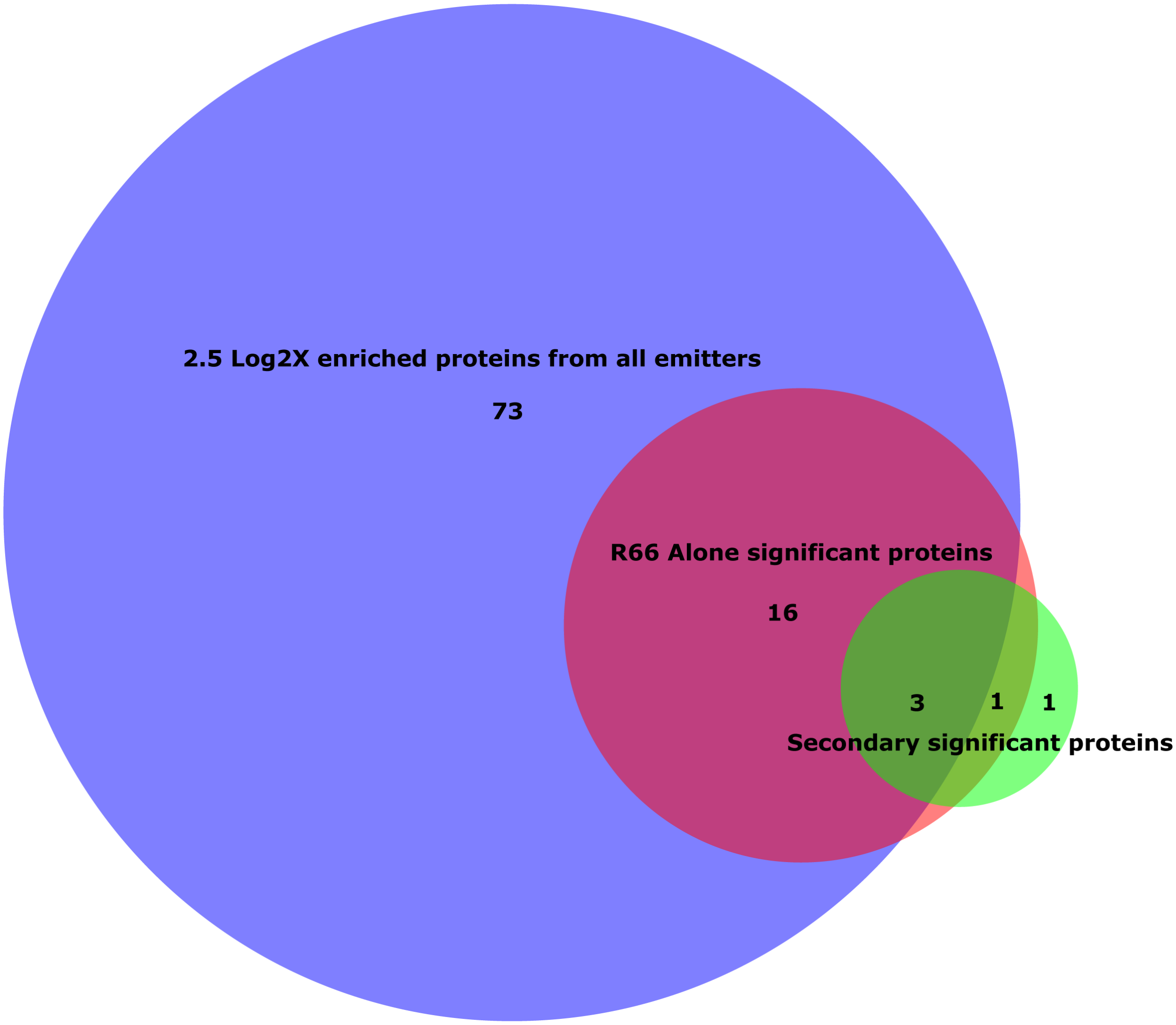


**SI Figure 5.** Comparing the labeling of R66$M_L$ alone and the secondary labeling strategy relative to the most enriched proteins from all eight Traz catalysts. The vast majority of the proteins identified by R66$M_L$ alone and most of the secondary emitter significantly enriched proteins are the same as those seen in the larger dataset.

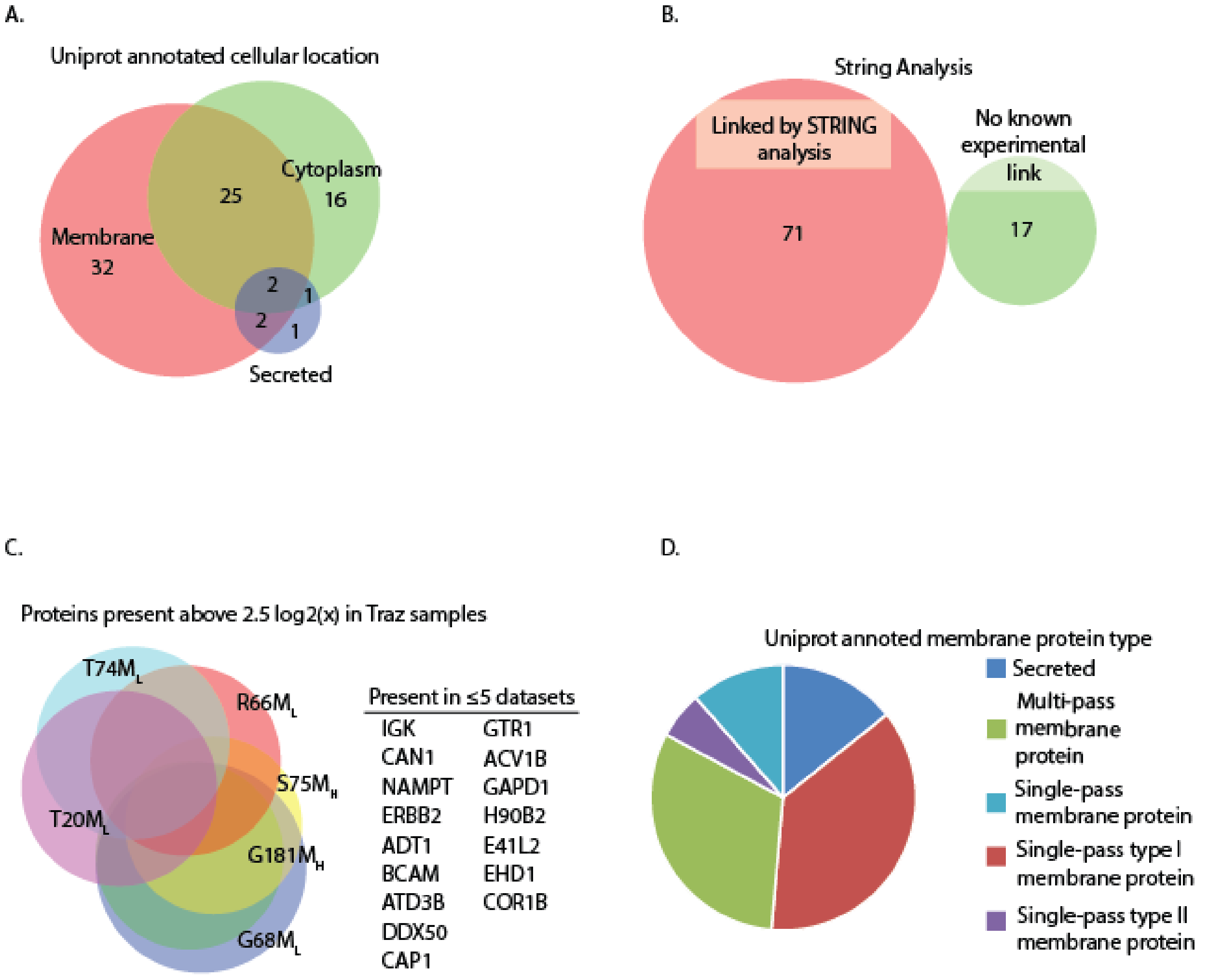


**SI Figure 6.** A. A Venn diagram showing the overlap for all high confidence identified proteins identified in the FA-emitter and the Fab-emitters. Virtually all of the high confidence proteins identified in the 8 Fab-emitters were detected in the FA-emitter experiment. B. A String analysis for eachof the highly enriched proteins from the Fab-emitter volcano plots. There was a strong association with HER2 as 71 of 88 proteins have a known link. C. A Venn diagram showing there is considerable overlap among highly enriched proteins from the Fab-emitters and a list of proteins present in at least four of the Fab-emitters. D. The cellular location GO annotation of the highly enriched proteins demonstrating that hits are predominantly identified as membrane or extracellular, with only 16 annotated as cytoplasmic

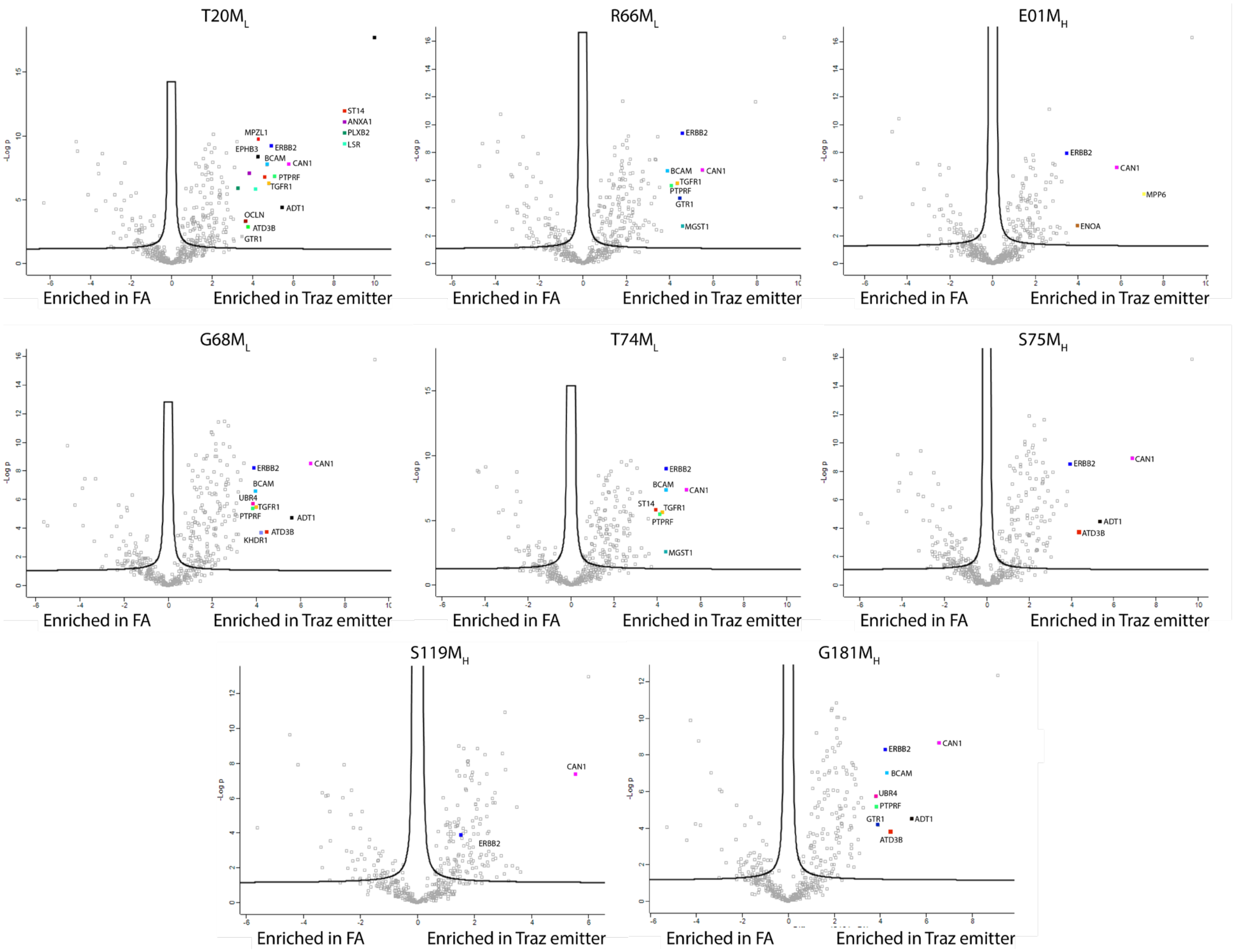


**SI Figure 7.** Volcano plots for each of the eight Traz catalysts used in the study. The most enriched and common proteins in each of the plots are labelled.

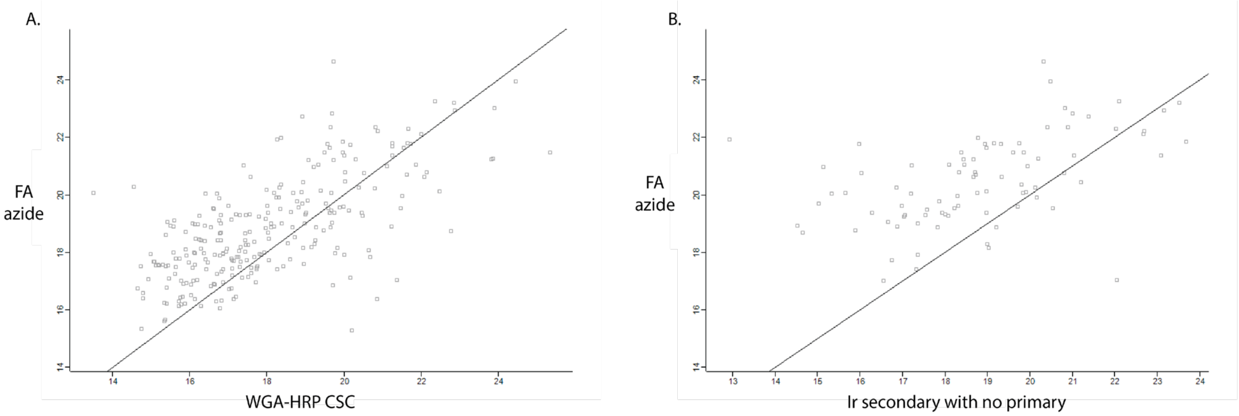


**SI Figure 8**. Comparing the LFQ values from CSC, FA azide, and unbound Ir secondary controls. A. The mean LFQ of the FA azide and WGA-HRP controls. The line is set to represent even LFQ values. B. The mean LFQ of the FA azide and Ir secondary without a primary control. The line is set to represent even LFQ values.

**SI Table 1.** Performance of the three controls chosen for comparison.

| Control | Proteins identified | Mean standard deviation (Log2x) |
|---|---|---|
| WGA-HRP | 543 | 0.39 |
| Fatty acid azide | 787 | 0.32 |
| Isotype | 150 | 0.67 |

**SI Table 2**. Mean LFQ values for experiments. Fab-catalyst sites are grouped together with means calculated for the 3 biological replicates and 2 technical replicates. The fatty acid catalyst experimental group is shortened to FA. The secondary-catalyst experiment is shortened as 2nd.

| PROTEIN IDS | G181$M_H$ | G68$M_L$ | S75$M_H$ | E01$M_H$ | R66$M_L$ | S119$M_H$ | T20$M_L$ | T74$M_L$ | FA | 2ND |
|---|---|---|---|---|---|---|---|---|---|---|
| **ENPL** | 22.806 | 22.871 | 22.868 | 23.269 | 22.752 | 22.916 | 22.657 | 23.030 | 22.843 | 21.419 |
| **PLEC** | 22.140 | 22.115 | 22.067 | 21.846 | 22.180 | 22.224 | 22.104 | 22.046 | 21.934 | 18.776 |
| **RB39A** | 21.386 | 20.331 | 21.379 | 23.222 | 21.933 | 21.935 | 23.198 | 23.254 | 20.853 | 19.253 |
| **HNRPF** | 22.234 | 22.367 | 22.256 | 21.986 | 21.884 | 22.325 | 21.949 | 21.866 | 21.806 | 20.309 |
| **EPIPL** | 22.470 | 22.437 | 22.616 | 20.774 | 21.695 | 21.616 | 21.907 | 21.373 | 20.982 | 19.237 |
| **HM13** | 20.227 | 21.157 | 21.131 | 21.660 | 21.127 | 20.663 | 21.034 | 21.417 | 20.739 | 19.771 |
| **RAB15** | 20.812 | 20.714 | 21.049 | 20.232 | 20.942 | 19.633 | 20.813 | 19.772 | 19.669 | 19.046 |
| **GBB2** | 19.917 | 20.817 | 20.768 | 20.788 | 20.840 | 19.837 | 20.734 | 20.955 | 20.036 | 16.945 |
| **PCAT1** | 19.884 | 20.810 | 20.840 | 20.031 | 20.636 | 20.575 | 20.884 | 20.536 | 20.232 | 15.645 |
| **PKP3** | 20.557 | 20.544 | 20.347 | 19.190 | 20.105 | 20.166 | 20.387 | 19.805 | 18.884 | 17.140 |
| **RP1BL** | 19.727 | 19.873 | 19.654 | 19.819 | 19.938 | 20.046 | 19.803 | 20.162 | 19.303 | 17.600 |
| **SURF4** | 19.822 | 20.523 | 19.992 | 18.448 | 19.918 | 20.562 | 19.902 | 20.304 | 19.442 | 14.597 |
| **ARF5** | 20.754 | 19.874 | 20.048 | 19.638 | 19.697 | 20.792 | 20.574 | 20.701 | 18.565 | 15.292 |
| **CTNB1** | 18.569 | 18.829 | 18.725 | 19.087 | 19.605 | 19.191 | 19.652 | 19.375 | 18.694 | 18.075 |
| **DESP** | 19.256 | 19.088 | 19.070 | 19.304 | 19.412 | 19.509 | 19.484 | 19.185 | 18.926 | 18.491 |
| **MIRO2** | 19.684 | 19.070 | 18.115 | 17.029 | 18.679 | 17.962 | 18.808 | 17.949 | 17.438 | 15.139 |
| **CAN2** | 18.776 | 18.915 | 18.880 | 17.853 | 18.644 | 18.616 | 18.744 | 18.146 | 16.929 | 14.718 |
| **DC1L1** | 18.147 | 17.650 | 17.276 | 17.635 | 18.442 | 17.926 | 18.726 | 18.376 | 16.909 | 16.101 |
| **L2GL2** | 17.516 | 17.388 | 16.431 | 17.393 | 18.429 | 18.562 | 18.112 | 16.799 | 17.423 | 14.676 |
| **AT1B3** | 18.827 | 20.515 | 19.717 | 18.680 | 18.424 | 18.657 | 18.969 | 19.842 | 17.401 | 18.795 |
| **G6PI** | 17.501 | 17.391 | 15.736 | 18.610 | 18.377 | 17.689 | 17.705 | 18.658 | 18.291 | 18.274 |
| **HMOX2** | 18.588 | 18.456 | 17.845 | 18.902 | 18.287 | 17.426 | 18.950 | 19.004 | 19.025 | 17.095 |
| **ASPH** | 19.465 | 19.594 | 18.799 | 17.726 | 18.214 | 18.365 | 18.240 | 17.851 | 18.605 | 15.045 |
| **RAB8B** | 16.833 | 18.185 | 19.090 | 17.309 | 18.057 | 17.910 | 17.793 | 18.720 | 18.699 | 17.508 |
| **PREX1** | 16.898 | 16.571 | 16.902 | 16.326 | 17.986 | 16.883 | 16.629 | 17.291 | 16.282 | 14.399 |
| **CNNM4** | 17.263 | 16.734 | 16.452 | 17.883 | 17.867 | 16.561 | 18.762 | 17.224 | 17.311 | 15.232 |
| **TLN2** | 17.953 | 18.362 | 17.663 | 18.303 | 17.814 | 18.376 | 18.087 | 18.398 | 16.894 | 16.313 |
| **SCRB1** | 15.679 | 15.709 | 15.884 | 17.752 | 17.812 | 15.626 | 18.695 | 17.587 | 18.695 | 22.586 |
| **ATPO** | 18.627 | 19.650 | 18.533 | 19.129 | 17.788 | 18.464 | 17.032 | 18.064 | 19.552 | 14.522 |
| **MOT4** | 18.664 | 17.916 | 15.667 | 18.083 | 17.729 | 18.810 | 18.523 | 16.975 | 17.193 | 14.626 |
| **NDUS7** | 17.443 | 19.044 | 17.745 | 15.950 | 17.695 | 17.595 | 18.407 | 16.226 | 16.779 | 14.637 |
| **CBPD** | 17.440 | 17.717 | 16.450 | 18.247 | 17.635 | 18.021 | 17.075 | 18.453 | 17.470 | 14.652 |
| **HIP1R** | 15.710 | 15.877 | 16.320 | 16.613 | 17.354 | 17.333 | 18.645 | 16.493 | 15.556 | 14.752 |
| **NDC1** | 17.345 | 15.532 | 16.339 | 16.024 | 17.325 | 16.630 | 16.223 | 16.504 | 15.915 | 14.648 |
| **FAD1** | 18.367 | 18.127 | 17.083 | 17.145 | 17.300 | 17.677 | 17.516 | 17.434 | 18.430 | 14.942 |
| **RAB31** | 16.741 | 16.258 | 15.759 | 16.644 | 17.265 | 17.075 | 17.871 | 17.093 | 17.180 | 15.077 |
| **SYMPK** | 15.983 | 17.022 | 16.282 | 16.079 | 17.235 | 16.815 | 16.769 | 16.524 | 16.840 | 14.517 |
| **ARFG2** | 16.404 | 16.294 | 15.861 | 17.200 | 17.234 | 16.178 | 17.243 | 16.962 | 16.130 | 14.268 |
| **KPBB** | 16.518 | 17.458 | 17.525 | 16.447 | 17.211 | 16.009 | 17.369 | 16.253 | 16.292 | 14.406 |

| | | | | | | | | | | |
|---|---|---|---|---|---|---|---|---|---|---|
| **RHEB** | 16.606 | 17.176 | 17.870 | 15.749 | 17.146 | 18.776 | 17.958 | 16.779 | 16.733 | 14.852 |
| **ITPK1** | 16.123 | 15.731 | 15.682 | 16.015 | 17.108 | 15.727 | 15.961 | 16.524 | 15.929 | 14.762 |
| **CUL1** | 16.831 | 16.829 | 15.954 | 17.270 | 17.079 | 17.247 | 16.862 | 17.538 | 16.464 | 14.657 |
| **SORT** | 16.450 | 17.385 | 17.678 | 17.210 | 17.038 | 17.097 | 17.224 | 17.543 | 16.107 | 14.273 |
| **GNAT3** | 17.373 | 16.739 | 16.053 | 16.468 | 17.031 | 15.680 | 16.898 | 16.457 | 15.955 | 14.707 |
| **ERLN2** | 17.019 | 17.859 | 16.563 | 17.034 | 17.026 | 16.545 | 17.935 | 16.633 | 15.860 | 14.438 |
| **DSG1** | 15.723 | 15.982 | 15.802 | 17.387 | 17.025 | 16.495 | 15.908 | 15.999 | 15.666 | 17.302 |
| **EXOC3** | 17.500 | 16.806 | 16.817 | 15.926 | 16.931 | 16.381 | 16.924 | 16.077 | 15.658 | 14.589 |
| **CUL3** | 16.179 | 16.687 | 16.545 | 17.813 | 16.901 | 16.560 | 15.885 | 17.599 | 15.834 | 14.753 |
| **AT2B2** | 17.689 | 17.498 | 16.601 | 17.817 | 16.838 | 17.658 | 17.542 | 18.415 | 16.659 | 15.526 |
| **OSBP1** | 17.091 | 17.879 | 17.515 | 16.801 | 16.807 | 17.252 | 16.031 | 16.869 | 15.984 | 14.433 |
